# Harnessing photosynthetic ATP for whole-cell biocatalysis in the cyanobacterium *Synechocystis*

**DOI:** 10.1101/2025.05.29.656799

**Authors:** Giovanni Loprete, Eleonora Traverso, Filippo Vascon, Marco Botteri, Marina Simona Robescu, Daniela Ubiali, Laura Cendron, Tomas Morosinotto, Elisabetta Bergantino

## Abstract

Photosynthetic organisms use sunlight to produce ATP and NADPH powering their metabolism. Harnessing these products for driving biocatalytic reactions would enable to develop clean and sustainable alternatives for chemical reactions. In this study, we present the first demonstration that ATP produced from the photosynthetic process, can fuel a biocatalytic transformation in the whole-cell configuration. This result was achieved by expressing in the cyanobacterium *Synechocystis* sp. PCC 6803 an ATP-dependent enzyme, the γ-Glutamyl-MethylAmide Synthetase from *Methylovorus mays* No. 9 (*Mm*GMAS). The expressed enzyme was able to drive in the transgenic strain the light-driven biosynthesis of L-theanine. Consumption of ATP by the recombinant *Mm*GMAS was even beneficial under strong illumination, protecting the photosynthetic electron transport from photodamage. These findings demonstrate the possibility of using photosynthetic microorganisms like *Synechocystis* as potential platform for sunlight driven biotransformations with wide potential biocatalytic applications. In this perspective, we further present the tridimensional structure of *Mm*GMAS, which explains its promiscuous in vivo activity and provides the basis for its rational evolution.

## INTRODUCTION

Cyanobacteria are photoautotrophic prokaryotes capable of converting CO_2_ into organic molecules using light energy through photosynthesis. They are responsible for the 20-30% of the earth’s organic carbon fixed and exhibit a photosynthetic efficiency that is potentially larger than that of plants (Waterbury et al., 1979; Li et al., 2008; Noreña-Caro and Benton 2018). Cyanobacteria are a potential green chassis for biotechnological applications, bringing together advantages from plants and bacterial worlds. While they perform photosynthesis as plants, yet their metabolic networks and molecular biology share similarities with bacteria, facilitating genetic manipulations (Rosgaard et al., 2012).

One major possibility to be explored is the use of cyanobacteria for biocatalysis performed in the whole-cell configuration, driving chemical reactions without the need of purifying enzymes and thus merging the sustainability and costs benefits. A crucial aspect to develop whole-cell biocatalytic processes is the supply of cofactors driving the desired chemical conversions. Oxygenic photosynthesis generates NADPH equivalents and reduced ferredoxin (Fd) from oxidation of water, thus representing an atom-efficient cofactor recycling system that avoids the requirement of suicide substrates and auxiliary enzymes. Consequently, researchers are currently focusing on utilizing *Synechocystis* sp. PCC6803 (*Synechocystis* from here on) to express heterologous NADPH-dependent enzymes to perform chemical reactions fueled by light (Park and Choi, 2017). Indeed, Baeyer-Villiger Monooxygenases (BVMOs) and Ene-Reductases (ERs) have already been employed in light-fueled whole-cell biotransformations to produce high-value molecules such as χ-caprolactone, ψ-valerolactone, levodione and *N*-methylsuccinimide (for a review see Malihan-Yap et al., 2022).

Besides oxidoreductases, ATP-dependent enzymes are indeed attractive biocatalysts due to their ability to drive challenging chemical reactions. Different in vitro ATP recycling systems have been developed such as acetate kinase-acetyl phosphate, pyruvate kinase-phosphoenolpyruvate, creatine kinase-creatine phosphate and polyphosphate kinase-polyphosphate. The application of these systems at scale is however inhibited by issues including (*i*) cost, stability and availability of the phosphate donor as well as (*ii*) requirement of multiple auxiliary enzymes to achieve AMP to ATP regeneration. (Tavanti et al., 2021)

In nature, the most efficient ATP production systems are found in the processes of cellular respiration and photosynthesis (Andexer et al., 2015). Photosystems produce ATP equivalents as well as NAPDPH, thus theoretically making *Synechocystis* a potential platform also for the expression of ATP-dependent enzymes. This possibility was explored here by expressing in *Synechocystis* a ψ-Glutamyl-MethylAmide Synthetase (GMAS, EC 6.3.4.12), an ATP-dependent enzyme that has raised an interest as a biocatalyst for the production of L-theanine (L-thea; 2-amino-4-(ethylcarbamoyl)-butyric acid; CAS 3081-61-6). L-thea is an unnatural amino acid contained in a wide range of nutraceutical formulations present on the market and used to increase the mental focus, while providing relaxation during task performances, to reduce stress and improve the quality of sleep (Matsumoto et al., 2005; Kimura et al., 2007; Yamada et al., 2008; Yoto et al., 2012; Hidese et al.,2019; Wang et al., 2021a). Currently, L-thea has been certified as a “Generally Recognized as Safe” (GRAS) ingredient by the Food and Drug Administration (FDA), and its demand is expected to grow significantly in the next few years.

The extraction of L-thea from its primary source, tea leaves (where it is present at about 7-21 mg/g of dry weight; Sharma et al., 2018) remains a complex and inefficient process. Chemical synthesis methods were developed but they are not economically competitive, while being laborious and risky, requiring numerous purification steps, and the use of hazardous chemicals (Ekanayake and Li, 2007; Sharma et al., 2018). An alternative and innovative approach for L-thea production is its biotechnological production using enzyme-catalyzed *in vivo* biotransformations. The literature reports the production of L-thea from sugar and ethylamine by fermentative processes employing a heavily engineered *E. coli* strain expressing a heterologous GMAS (Fan et al., 2020), or the two bacteria *Corynebacterium glutamicum* (Ma et al., 2020; Yang et al., 2024) or *Pseudomonas putida* KT2440 (Benninghaus et al., 2021).

In this work, we expressed in *Synechocystis* the GMAS from the methylotrophic bacteria *Methylovorus mays* No. 9 (*Mm*GMAS; Yamamoto et al., 2008), which catalyzes the condensation of L-glutamic acid and ethylamine in the presence of molar equivalents of ATP as co-substrate (Fig. 1). The expressed enzyme was functional as a biocatalyst in whole-cell configuration, driving light-fueled biotransformation of L-thea. Under strong illumination, the consumption of ATP by the recombinant enzyme increased the photosynthetic electron transport capacity. We finally describe the structure of the enzyme, which helps explaining its promiscuous catalytic activity in the cyanobacterial cell, opening perspective for further engineering aimed at optimizing its enzymatic activity.

**Fig. 1.**
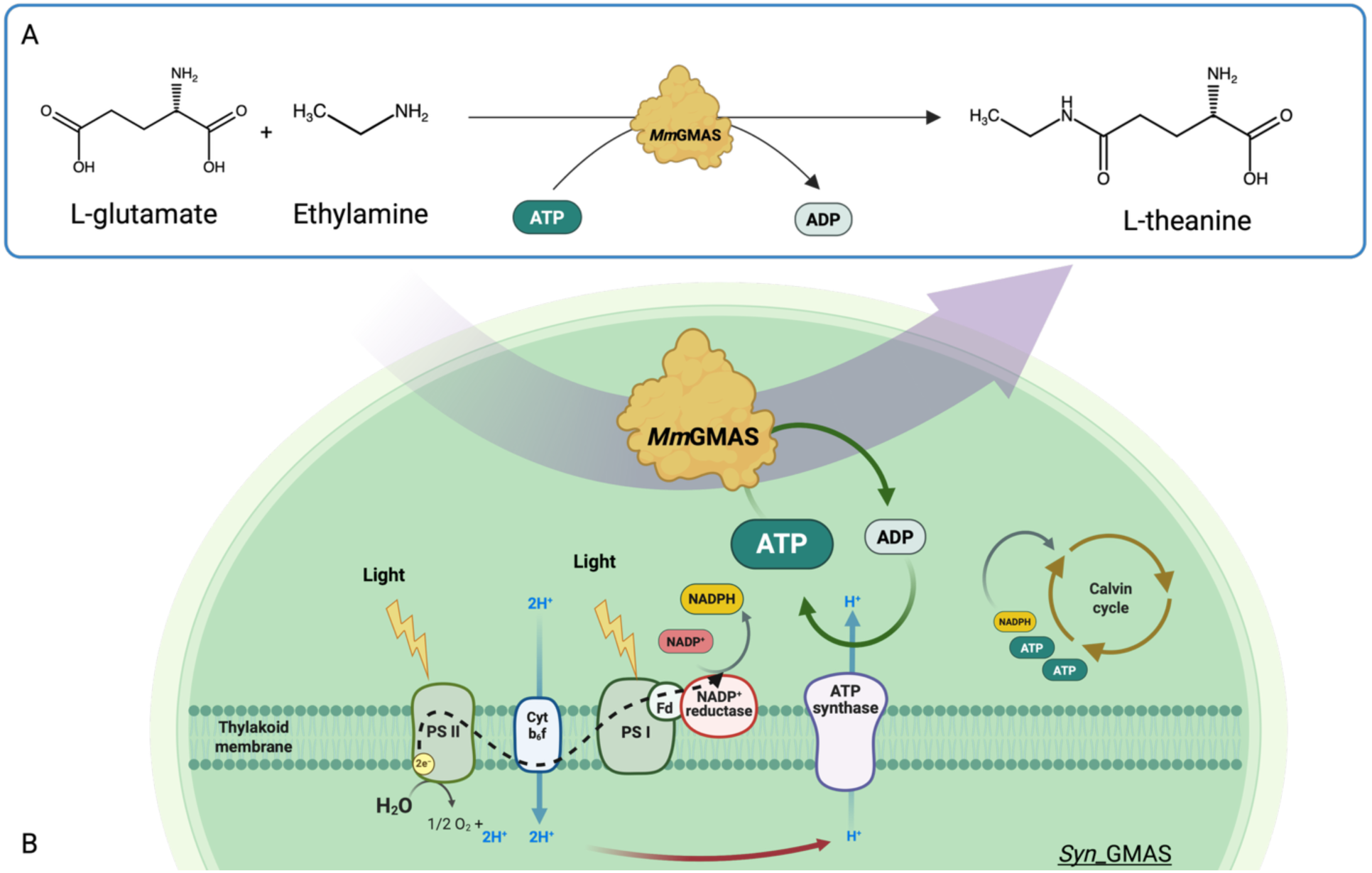
(A) In vitro biocatalytic reaction producing L-thea, catalyzed by recombinant *Mm*GMAS. **(B)** Sketch of the whole-cell biotransformation investigated in this study and performed by the *Mm*GMAS-expressing transgenic strain *Syn*_GMAS. Fluxes of electrons, NADPH and ATP cofactors, and protons (H^+^) are represented by dotted black, grey, green and red arrows, respectively.

## RESULTS and DISCUSSION

### A transgenic *Synechocystis* strain expressing *Mm*GMAS produces L-theanine from L-glutamic acid and ethylamine in whole-cell biotransformation

A homoplasmic, stable *Synechocystis* strain expressing the enzyme *Mm*GMAS under the control of the endogenous strong promoter P_cpc560_ was generated (Fig. S1; Loprete et al., 2025). The cyanobacterial strain, *Syn*_GMAS, was validated for the presence of the heterologous gene and *Mm*GMAS expression was assessed by western blotting (Fig. S2, S3). A transgenic strain resistant to kanamycin but lacking the coding sequence for the enzyme (*Syn*_UV) was also produced and used as control throughout the whole work (see Supplementary Information).

In vivo biotransformations were set-up to test the activity of *Mm*GMAS, which also requires (*i*) membrane permeation by the substrates provided in the growth medium and (*ii*) product secretion. Substrates – ethylamine and L-glutamic acid (L-glu) – were added to the culture medium at a final concentration of 1 mM at a working OD_730_ = 5. L-Theanine formation over time was monitored by analyzing the supernatant by Thin Layer Chromatography (TLC; Fig. S4A and S4B) and showed that L-thea was present in the medium collected from the *Syn*_GMAS culture. Identity of the product was also confirmed by analyzing the culture supernatant by ElectroSpray Ionization-Mass Spectrometry (ESI-MS, Fig. S5). These results demonstrated that the heterologous *Mm*GMAS was active and also that both substrates and product can permeate the cellular membranes.

Since L-glu is a natural amino acid, we further tested whether *Syn_*GMAS could synthesize L-thea by providing only ethylamine, relying on endogenous L-glu. The results of such biotransformation assays, shown in Fig. S4C, demonstrated that L-glu was dispensable.

### Yields and kinetics of the biotransformation producing L-theanine

Once validated, reaction kinetics were assessed using varying substrate concentrations. Aliquots of the supernatant from the biotransformations were collected at defined time points, derivatized and analyzed by Reverse Phase-HPLC.

The samples containing 5 mM and 10 mM of both substrates showed comparable levels of L-thea production and kinetic parameters (Table 1), whereas at 2.5 mM the production is reduced by approximately a half.

**Tab. 1.** On the left, measurements of rates and specific activities of *Syn*_GMAS biotransformations presented in Fig. 2A. Calculations were made in the interval of 0-24 hours from the addition of the substrates, corresponding to the observed maximum production period. On the right, measurements of rates and specific activities of *Syn*_GMAS biotransformations presented in Fig. 2B. Calculations were made in the interval of 4-6 hours from the addition of ethylamine, corresponding to the observed maximum production. Data are the average of three independent replicates.

| L-glutamate and ethylamine (mM) | Rate (mM/h) | Specific activity (U/g<sub>DCW</sub>) | Ethylamine (mM) | Rate (mM/h) | Specific activity (U/g<sub>DCW</sub>) |
| --- | --- | --- | --- | --- | --- |
| 2.5 | 0.021±0.002 | 0.27±0.02 | 1 | 0.021±0.001 | 0.28±0.01 |
| 5 | 0.047±0.003 | 0.66±0.04 | 2.5 | 0.041±0.001 | 0.57±0.02 |
| 10 | 0.046±0.002 | 0.64±0.02 |  |  |  |

The consumption over time of substrates L-glu and ethylamine showed that while the latter was constantly consumed since its addition, the concentration of the former remained nearly constant up to 24 hours suggesting that that L-thea was initially produced by consuming endogenous L-glu Fig. 2A. Mass balance related to L-glu, reported in Fig. S6, increased over time up to 24 hours and then started to decrease, thus confirming that endogenous L-glu was initially consumed. Mass balance related to ethylamine, showing a constant decrease, confirmed that it was steadily consumed all along the biotransformation.

**Fig. 2.**
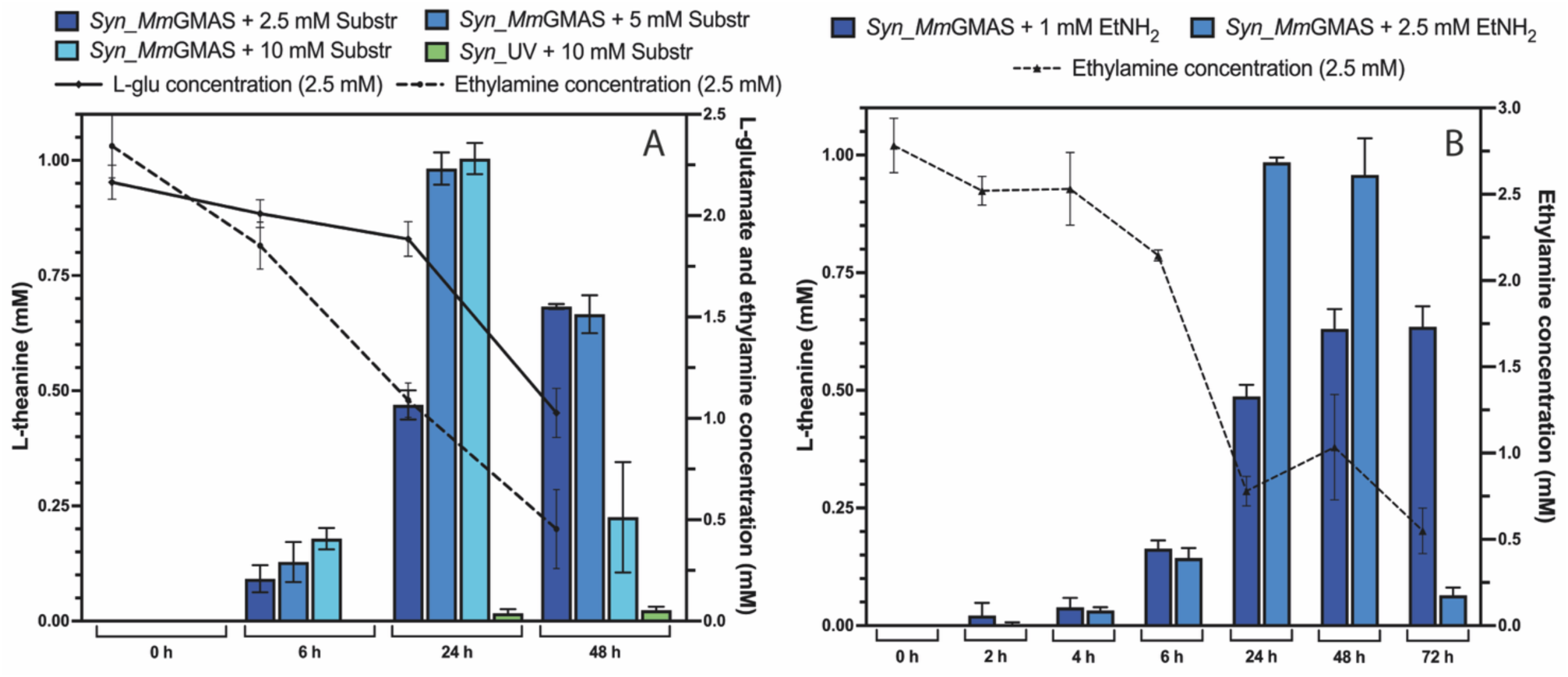
(**A**) Results of RP-HPLC analysis of *Syn*_GMAS whole-cell biotransformations supplemented with different concentrations of L-glutamate and ethylamine (2.5, 5 and 10 mM) at different times. Concentration of L-glu and ethylamine over time are represented by the continuous and dashed black lines respectively. Control strain (*Syn*_UV) biotransformation was performed in the presence of 10 mM substrates. (**B**) Results of RP-HPLC analysis of *Syn*_GMAS whole-cell biotransformations supplemented with different concentrations of ethylamine (1 and 2.5 mM) at different times. Concentration of ethylamine over time is represented by the dashed black line.

As before, biotransformations performed by adding only ethylamine confirmed the feasibility of L-thea production relying on endogenous L-glu (Fig. 2B). Moreover, starting from 2.5 mM ethylamine, L-thea production peaked at 1 mM, matching the highest yield obtained when providing both substrates in the culture medium at 5 mM and 10 mM. Additionally, rates and specific activity, reported in Tab. 1, are similar to the ones calculated in the presence of both substrates.

The L-thea content was also observed to decrease after 24 hours from the beginning of the biotransformations at 5 and 10 mM L-glu and ethylamine (Fig. 2A). This could be due to L-thea instability in the medium or because of reverse reaction by *Mm*GMAS. Both hypotheses were excluded by in vitro tests (Fig. S7, S8A and S8B). Notably, purified recombinant *Mm*GMAS was fully proficient in L-thea synthesis, while no degradation was detectable.

A third possibility was that L-thea could be metabolized by the cyanobacterial cells. This was tested by cultivating *Syn*_GMAS and the control strain in the presence of exogenously added L-thea. Both strains exhibited L-thea consumption, with equivalent rates (0.016 ± 0.006 and 0.017 ± 0.013 mM/h for *Syn*_UV and *Syn*_GMAS respectively), suggesting L-thea is metabolized by endogenous enzymes of *Synechocystis*.

L-thea might serve as a nitrogen source in *Synechocystis*, in line with what reported for nitrogen storage and transport in tea plants (Tsushida and Takeo, 1985; Li et al., 2020; Chen et al., 2021), where it is metabolized by glutaminases like CsPDX2.1 that was shown to be active on L-thea (Fu et al., 2020). At least two *Synechocystis* enzymes with glutaminase activity have been reported but never tested with L-thea (Ravasio et al., 2002; Dossena et al., 2007; Zhou et al., 2008). To test if L-thea could be hydrolyzed and employed as an alternative nitrogen source in *Synechocystis* as well, we monitored the growth of the wild-type strain – unable to fix nitrogen – in a medium devoid of nitrogen, in the presence or absence of added L-thea. Without L-thea, wild-type *Synechocystis* rapidly showed the characteristic nitrogen-starvation phenotype (Li and Sherman, 2002; shown in Fig. S9A), including the reduction of chlorophyll content (Fig. S9B). Conversely, L-thea supplementation prevented the nitrogen-starvation and chlorophyll content reduction (Fig. S9). These evidences confirmed that L-thea can be metabolized and used as a nitrogen source by the cyanobacterium.

We can conclude that in biotransformations employing *Syn*_GMAS (*i*) the highest production of L-thea (174 mg per liter of culture, 1 mM) was reached at 24 hours from the addition of both substrates, L-glu and ethylamine; (*ii*) in that time span, the production could be sustained by the endogenous reserve of L-glu alone; (*iii*) all along the process, produced L-thea is in equilibrium between the medium and the cells, where it is hydrolyzed by endogenous enzyme(s).

### *Mm*GMAS affects the cyanobacterial metabolism, leading to a decreased exponential growth rate

Once the feasibility of the biotransformation was verified, we investigated the impact of the expressed heterologous protein on *Synechocystis* growth. Fig. 3A shows that, in the absence of added substrates, *Syn*_GMAS and control *Syn*_UV cultures showed similar growth and biomass accumulation (1.23 ± 0.16 g_DCW_/L vs. 1.30 ± 0.15 g_DCW_/L respectively). However, *Syn*_GMAS showed a small delay in the exponential phase of growth. This could be ascribed either to the metabolic burden due to *Mm*GMAS (over)expression, or to its basal activity on one or more endogenous molecules thus consuming ATP. The enzyme has in fact been reported to be a promiscuous catalyst, capable of producing many different γ-glutamyl compounds from ammonia up to more sterically hindered amines, such as tryptamine or phenylethylamine (Yamamoto et al., 2008; Pan et al., 2020). Considering the presence of various endogenous amines in *Synechocystis*, *e.g.* ammonia, (see Fig. S10), a basal activity by the expressed *Mm*GMAS is likely.

**Fig. 3.**
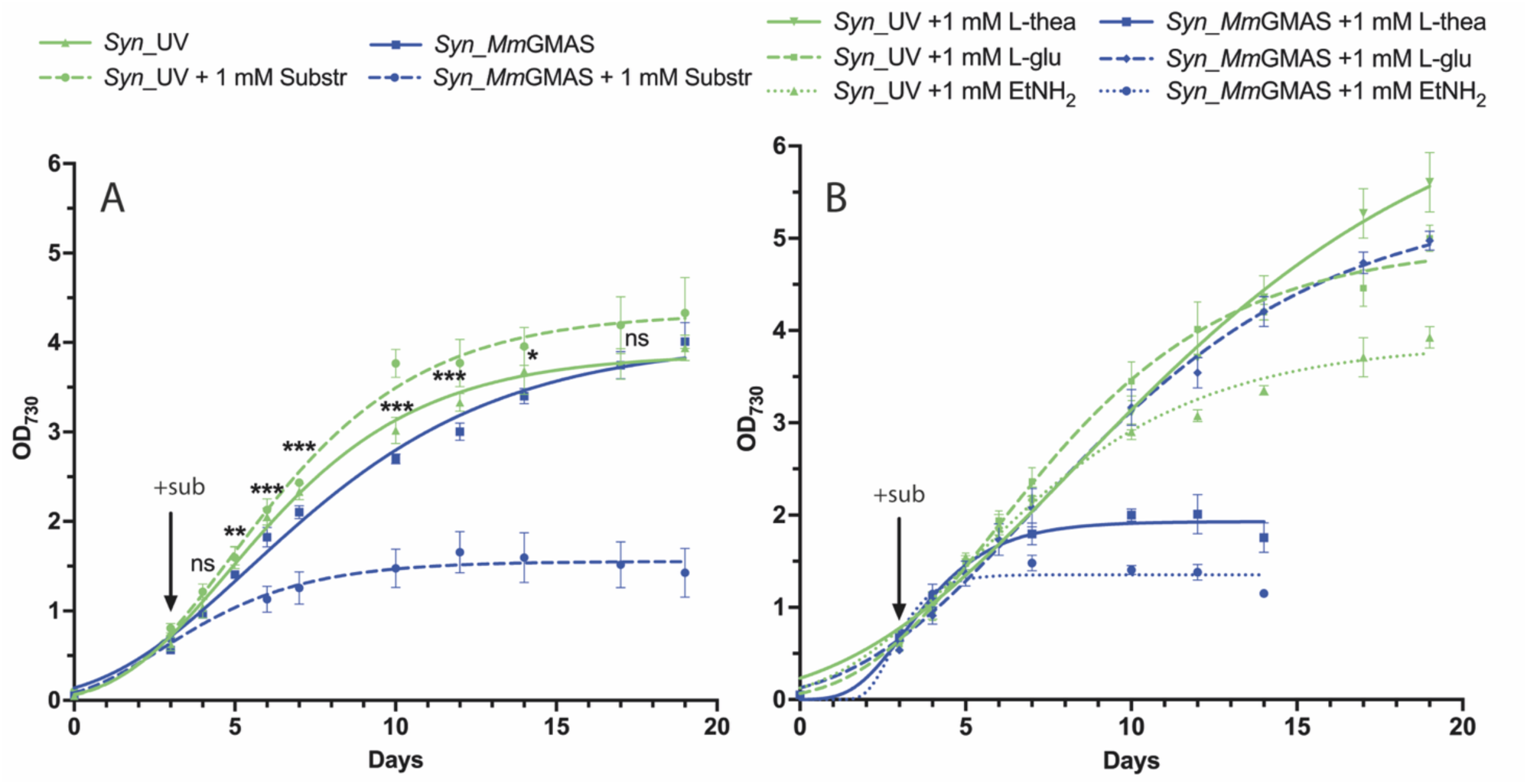
Growth curves. (**A**) Curves comparing growth rates of Syn_GMAS and Syn_UV strains in the absence and presence of 1 mM L-glutamate and ethylamine. Statistical significance (two-sample t-test) is represented by ns (not significant) and asterisks (significant, *p<0.05, **p<0.01, ***p<0.001) and refers to each time-point measure in the absence of substrates. (**B**) Curves comparing growth of strain Syn_GMAS and Syn_UV in the presence of 1 mM L-glutamate, L-theanine and ethylamine separately. Cultures (three independent biological replicates for each strain and each condition) were cultivated in flasks under continuous shaking and in standard cultivation conditions.

When substrates were added to the medium, however, the growth profile of the two strains significantly changed and while the presence of both L-glu and ethylamine are well tolerated by the control, their addition drastically inhibited *Syn*_GMAS growth (Fig. 3A).

Individual substrates or products were added separately to both *Syn*_UV and *Syn*_GMAS (Fig. 3B). Ethylamine did not impact *Syn*_UV growth, while L-glu and L-thea had an evident beneficial effect (Fig. 3B). On the contrary, *Syn*_GMAS responded positively only to the addition of L-glu, while its viability was compromised by the presence of either ethylamine or L-thea suggesting a growth inhibition observed associated with the heterologous enzyme’s activity when ethylamine is present, added or produced by L-thea hydrolysis.

### *Syn*_GMAS is characterized by a decreased endogenous free ATP content and an enhanced photosynthetic electron transport rate

In cyanobacteria the light-excited electrons drive the reduction of NADP⁺ into NADPH and contribute to the establishment of a proton gradient, then exploited by the ATP synthase to produce ATP, sustaining cellular growth. We thus monitored the impact of the ATP consuming enzyme on endogenous ATP levels and photosynthetic efficiency.

Intracellular ATP results 10% lower in *Syn*_GMAS compared to *Syn*_UV (Fig. 4A) which could explain the reduction in exponential growth rate observed for *Syn*_GMAS. In biotransformations, two hours after the addition of the substrates, the ATP consumption is strongly enhanced (Fig.4B), further confirming the enzyme activity and also explaining the growth inhibition of *Syn*_GMAS in the presence of ethylamine (with or w/o L-glu, Fig. 3A and 3B). These results clearly confirm that the enzyme’s activity was ATP dependent.

**Fig. 4.**
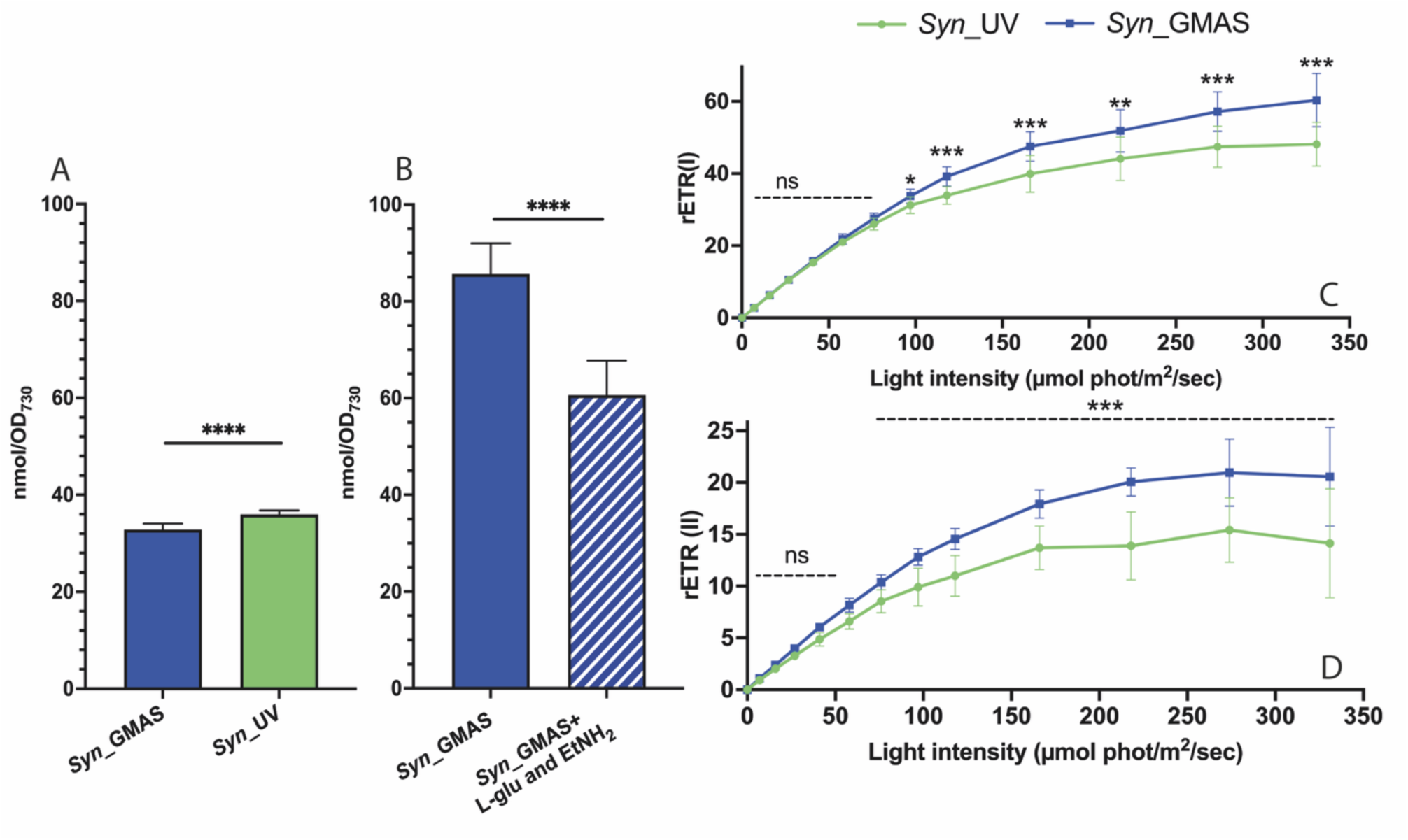
ATP content and Electron Transport Rates of PSII and PSI (rETR). (**A**) ATP content of *Syn*_GMAS and *Syn*_UV grown in standard conditions (up to exponential phase, BG11 and 50 μmol photons •m^-2^ •s^-1^) in the absence of exogenous substrates. (**B**) ATP quantification of *Syn*_GMAS whole-cell biotransformations after 2 hours of cultivation in the presence or absence of 1 mM substrates, in BG11 and under 150 μmol photons •m^-2^ •s^-1^ constant illumination. (**C**) Measurements of relative Electron Transport Rate of PSI and PSII, rETR(I) and (**D**) rETR(II) respectively, in *Syn*_GMAS and control *Syn*_UV strains. Statistical significance (two-sample t-test) is represented by ns (not significant) and asterisks (significant, *p<0.05, **p<0.01, ***p<0.001 and **** p<0.0001)

Maximal photosynthetic efficiency (Fv/Fm) was evaluated in dark adapted *Syn*_UV and *Syn*_GMAS strains grown in the absence of any substrate and showed no significant differences (0.30 ± 0.04 and 0.31 ± 0.06 respectively). When exposed to increasing light intensity, the relative PSII electron transport rate (rETRII) was higher for *Syn*_GMAS than for the control (Fig 4D).

Similarly, PSI-associated electron transport capacity (rETRI) was higher in *Syn*_GMAS when exposed to higher light intensities(Fig. 4C). This increase can be ascribed solely to the presence of the heterologous enzyme consuming ATP. Thanks to this activity, ATP consumption is increased, stimulating a higher electron transport rate. This reduces the saturation of photosynthetic ETR at higher intensities which can be beneficial to protect from photodamage.

### Structural characterization of *Mm*GMAS

The above presented data suggested a promiscuous behavior in vivo of *Mm*GMAS. The determination of its tridimensional structure was thus pursued to gain deeper insights into its catalytic activity and potential for optimization in biocatalytic applications, since the only experimental structure belonging to this enzymatic class is the one of the more specific *Rh*GMAS (PDB: 7CQL, 7CQN, 7CQQ, 7CQU, 7CQW and 7CQX; Wang et al., 2021b).

*Mm*GMAS was recombinantly produced and purified from *E. coli* cultures (Figs. S11A, S11B, S12); we obtained the crystal structure of the complex *Mm*GMAS-ATPγS at 2.65 Å (PDB 9QUR; Tab. S3), showing that the enzyme is organized as a homo-dodecamer composed of 2 stacked hexameric rings of *Mm*GMAS subunits (Fig. 5A and 5B). Details of interactions among protein chains are illustrated in the Supplementary Results section and Fig. S13 (Supporting Information).

**Fig. 5.**
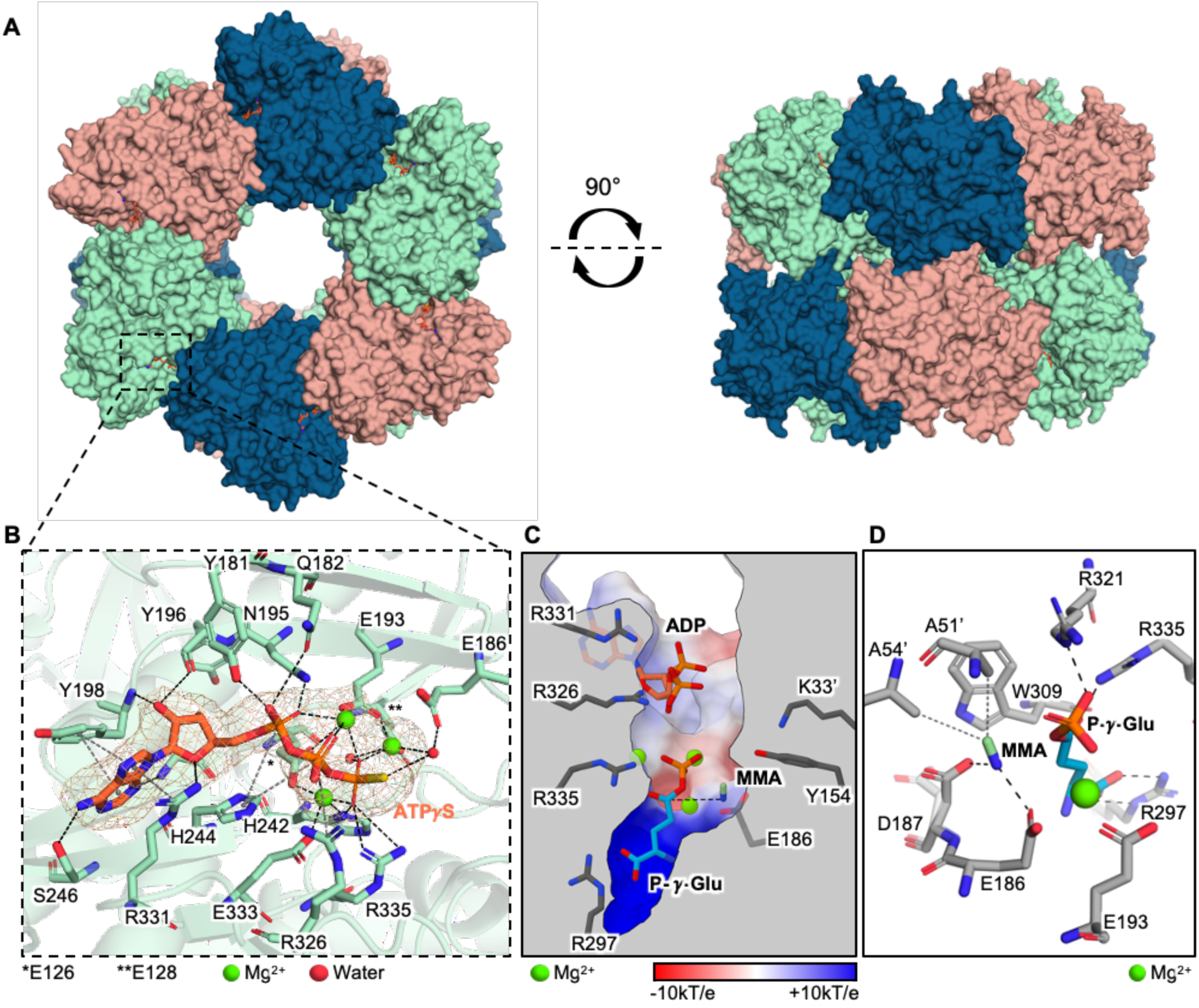
Crystal structure and computational modelling of MmGMAS. (**A**) Front and top view of the *Mm*GMAS homododecamer (PDB 9QUR). (**B**) Interactions between ATPγS, Mg^2+^ and *Mm*GMAS residues. Omit map of ATPγS and Mg^2+^ cations at 2.5 σ is displayed. H-bonds and metal coordination bonds are displayed as black dashed lines; salt bridges, π-stacking and cation-π interactions are shown as gray dashed lines; hydrophobic interactions are depicted as grey dotted lines. (**C**) In silico generated model of the quaternary complex *Mm*GMAS/ADP/P-γ-Glu/MMA. Color gradient depicts the surface electrostatic potential of *Mm*GMAS active site. The dashed line represents the nucleophilic attack of MMA amine group on P-γ-Glu Cδ. (**D**) Interactions between *Mm*GMAS residues, MMA and P-γ-Glu. Hydrophobic interactions are depicted as grey dotted lines while electrostatic contacts are shown as black dashed lines.

*Mm*GMAS shares an identity of 44% (59% homology) with the primary sequence of *Rh*GMAS; high conservation of the tridimensional fold and residues of the active site is also observed. Most relevant structural differences are located on flexible secondary structures which, despite not directly involved in *Mm*GMAS catalytic cleft, are located both upstream and downstream substrates and cofactors binding sites (the β-hairpin 132-143 and the random-coil loops 248-252 and 256-267), and might impact catalytic activity, protein stability and dynamics by epistatic effects. These observations support the hypothesis that the substrate specificity of wild-type *Mm*GMAS relies on the plasticity of the catalytic site and the hexamer assembly, likely explaining its reported higher promiscuity with respect to *Rh*GMAS (Pan et al., 2020; Wang et al., 2021b).

To better understand substrate binding in *Mm*GMAS, a quaternary complex model including ADP, monomethylamine (MMA), and L-γ-glutamyl phosphate (P-γ-Glu) was predicted. In comparison to the previously proposed model, our one showed MMA occupying a novel binding site closer to the Cδ of P-γ-Glu, positioning it effectively for nucleophilic attack promoting the addition of the amine (Fig. 5B and Supporting Information). This binding is stabilized by hydrophobic and electrostatic interactions involving the key residues Ala51’ and Ala54’ (Fig. 5D). Moreover, molecular dynamics simulations confirmed a higher stability of MMA in this new site compared to the previous model (Supporting Information and Fig. S14). These findings are supported by prior mutagenesis studies, particularly the essential role of Glu186 and Asp187 (Glu179 and Asp180 in *Rh*GMAS; Fig. 5D; Wang et al., 2021b), highlighting the functional relevance of the proposed binding mode for *Mm*GMAS. Beside explaining the observed promiscuity in *Synechocystis*, *Mm*GMAS experimental X-ray structure together with these modeling and dynamics studies furnish valuable information for further engineering and evolution of the biocatalyst for both in vivo and in vitro applications (Fan et al., 2024).

## CONCLUSIONS

This study represents the first demonstration of the feasibility of an approach of whole-cell biotransformation that exploits ATP generated through photosynthesis. Our *Synechocystis* strain expressing *Mm*GMAS produces L-theanine in a light-fueled reaction.

As commonly observed with recombinant enzymes expressed in active form within host cells, *Mm*GMAS was found to affect cellular metabolism by consuming endogenous ATP, thereby altering metabolic equilibria and leading to reduced growth rates. Notably, however, the cyanobacterial photosystems appear to be capable of sustaining the ATP-dependent activity of *Mm*GMAS for the time required to achieve the highest bioconversion.

The selected enzyme and the specific catalyzed reaction were instrumental to establish ATP as a viable and tunable energy currency for driving biocatalysis in *Synechocystis*. This represents a breakthrough in the valorization of this organism as promising platform to exploit sunlight and CO_2_ to sustainably produce high value chemicals. Fundamental research and genetic manipulation — focused on enlarging molecular toolkits, enhancing light-harvesting capabilities, optimizing the operation of photosystems (PSs), minimizing parallel side reactions, and enabling structure-driven rational evolution of the enzyme(s)— are still required to address factors that limit the transition to industrially relevant applications, in a profitable, fair competition with well-established heterotrophic systems (Schmelling and Bross, 2024).

## METHODS

### Enzymes and reagents

Polymerases, restriction enzymes, DNA oligonucleotides (Table S1), standard L-theanine and L-glutamate were purchased at Thermo Fisher Scientific (United States). *Mm*GMAS coding sequence was purchased at GeneArt, Thermo Fisher Scientific (United States). Standard ethylamine and other reagents were purchased at Merck (Merck Life Sciences, Germany).

### Bacterial strains, cloning and Synechocystis transformation

The following strains were used: *Synechocystis* sp. PCC 6803 purchased at Pasteur Culture collection of Cyanobacteria (PCC, France), *E. coli* DH-5a and BL21 (DE3) at New England Biolab (United States). Cloning in *E. coli* and PCR amplifications were performed by routine methodologies. Plasmid pSuperP_*Mm*GMAS (Fig. S1) was constructed starting from the *Synechocystis* empty plasmid pSuperP_UV (Loprete et al., 2025). NcoI and NotI restriction sites at the 5’ and 3’ termini respectively were used to clone *Mm*GMAS coding sequence fused to a N-Terminus 6XHisTag. The protocol for *Synechocystis* transformation, using plasmid DNAs digested by DraI, was the one based on phosphate deprivation described by Pope et al. (2020).

### *Synechocystis* standard cultivation conditions, Optical Density measurements and dry cell weight

Standard culture cultivation was performed in BG11 medium (Tab. S2) continuous shaking (150 rpm) and under constant light illumination, 50 μmol photons •m^-2^ •s^-1^. Cultivation flasks and Corex tubes used for biotransformations were covered with a hydrophobic cotton cap allowing air exchange while limiting medium evaporation. Cyanobacterial population density was estimated from the turbidity of the culture, typically expressed as Optical Density at 730 nm (OD_730_). Dry cell weight was measured by completely drying 10 mL of cyanobacterial cultures on 0.2 µM nylon filters, which were dried overnight at 60 °C.

### Total protein extraction from *Synechocystis* cells and Western Blot analysis

Expression of the *Mm*GMAS in *Syn*_GMAS was verified by Western blotting as already described in Loprete et al. (2025). Briefly, total protein extract was obtained by harvesting cell cultures in exponential growth phase by centrifugation, washed, and resuspended in resuspension Buffer (50 mM HEPES-NaOH pH 7.5, 30 mM CaCl_2_, 800 mM sorbitol, 1 mM ε-amino-n-caproic acid), then homogenized using One Shot Cell disruptors (Costant Systems, United Kingdom). After centrifugation, soluble protein content was quantified by Bradford assay (SERVA Electrophoresis GmbH, Germany) and 10 µg were run in 12% UREA-PAGE. Western Blot was performed using primary Mouse monoclonal anti His-Tag antibody HRP-conjugated (SB194b, Southern Blotting, USA). VWR® Imager CHEMI Premium was used for chemiluminescence detection (VWR International s.r.l., Italy).

### Chlorophylls extraction and quantification

*Synechocystis* wild-type was cultivated in modified BG11 (nitrogen depleted medium, Tab. S2) in the presence and absence of L-theanine. After 48 hours, cultures were diluted by a factor of 10 to estimate the cellular content. Samples (20 µL) were deposited in the cell counting chamber, allowed to settle and counted using the Cellometer Auto X4 Cell Counter (Nexcelom Bioscience, United States). Counts were performed three times on three independent replicates. In parallel 2 mL of each replicate were harvested 20 min at 6000xg. After removal of the supernatant, cells were resuspended in *N,N*-dimethylformamide (DMF) and left overnight at 4 °C in darkness to extract chlorophylls. Then, the solutions were harvested 15 min at 6000xg to separate cellular debris and quantified using a Cary60 UV-Vis spectrophotometer (Agilent, United States).

### Spectroscopic analyses

Fluorescence and P700 measurements were carried out with Dual-PAM-100 fluorometer (Walz-Germany) on samples grown up to early exponential phase (OD_730_=1.5-2) under constant light illumination of 60 µmol photons m^2^ s^-1^. Prior to the analyses, samples were dark-incubated for 4 minutes and measurements were performed on a cell suspension of OD_730_=2 with sample loaded in the 1-cm rectangular quartz cuvette according to Patil et al., 2020.

Samples were exposed to increasing intensities of actinic red light and PSII and PSI-related parameters were calculated upon saturating pulses of 5,000 µmol photons m^2^ s^-1^, 600 msec (as in Patil et al., 2020; Zhou et al., 2016). Fv/Fm = (Fm−F0)/Fm was determined as the maximum quantum efficiency of PSII in the dark-adapted state. Relative electron transport rate of PSII and PSI (respectively as rETRII and rETRI) were estimated from PSII and PSI yield (Y(II) and Y(I)) calculated respectively as Fv’/Fm’ = (Fm’−F0)/Fm’ and 1-Y(ND)-Y(NA) (where Y(ND)= P-P0/Pm and Y(NA)= Pm-Pm’/Pm according to Klughammer and Schreiber, 2008).

### Intracellular free ATP quantification

Intracellular free ATP was measured by the ATP Determination kit (A22066, Thermo Fisher Scientific, United States). *Syn*_GMAS and *Syn*_UV strains were grown in continuous shaking at 150 rpm and in standard cultivation conditions up to early exponential growth phase (OD_730_=1.5-2). Aliquots of samples (150 µL) were immediately frozen in liquid nitrogen. After addition of an equal volume of glass beads (150-212 um, Sigma G-1145), homogenization was performed using the Bullet Blender Storm Pro cell homogenizer (Next Advance, United States). Samples were then centrifuged 1 min at 20000xg at 4 °C. ATP quantification was performed according to the manufacturer’s instructions.

### TLC and ESI-MS

Thin-Layer Chromatography (TLC) silica glass plates 60G F_254_ (Merck Life Sciences, Germany) were used. The mobile phase was set on the basis of literature and experimental trials. Finally, the eluent NH_3_/EtOH (70/30) for L-glutamate, ethylamine and L-theanine separation was used. Detection was performed using 0.1% w/ v Ninhydrin in EtOH under heat.

For mass analysis supernatant samples collected (2 mL) were dried under N_2_ flow and then suspended in MeOH (300 µL). Subsequently, the mixture was centrifuged to eliminate insoluble residues and then diluted 1:10 in MeOH for direct MS injection and analysis. MS detection was performed by using a linear ion trap mass spectrometer (LTQ) equipped with an electrospray ion source (ESI) (Thermo Scientific, San Jose, CA, USA) and controlled by X-calibur software (2.0.7 version). The following MS parameters were applied: positive ion mode, scan range 150–1000 m/z in full scan mode, source voltage 4.6 kV, capillary voltage 30 V, sheath gas flow rate 10 (arbitrary units), auxiliary gas flow rate 4 (arbitrary units), capillary temperature 250 °C, and tube lens voltage 80 V.

### Supernatant derivatization and RP-HPLC analysis protocol

Supernatant samples for RP-HPLC were isolated after centrifugation of culture samples at 13000xg for 15 minutes. 120 µL of methanol were added to 60 µL of supernatant. After 25 minutes centrifugation at 13000xg, samples were derivatized before RP-HPLC analysis. The protocol was optimized starting from the one described in Perucho et al., 2015. Derivatization was performed mixing 150 µL of samples with 75 µL of derivatization mix (32 mg of *O*-phthaldialdehyde dissolved in 800 µL of methanol, 7140 µL of 0.4 M borate buffer pH 9.5 and 60 µL of 3-mercaptopropionic acid). After 5 minutes at room temperature, the derivatization was quenched by the addition of 37.5 µL 5% v/v acetic acid and then immediately analyzed. RP-HPLC analyses were carried out on a Nucleosil 100 C-18 column (25 cm x 4.6 cm, 5 μm) (Altmann Analytik, Germany), employing as mobile phase A: 5% methanol/95% sodium acetate 0.05 M pH 5.88 and mobile phase B: 70% methanol in water. After injection (20 μL) in the Beckman Gold HPLC system (Beckman coulter, United States), the analysis was performed by a gradient elution as follows: 0-13 minutes (A 75%-B 25%-> A 0%-B 100%), 14-15 minutes (A 0%-B 100%), 16-20 minutes (A 0%-B 100%-> A 75%-B 25%), 21-25 minutes (A 75%-B 25%).

### Protein overexpression and purification

The coding sequence of *Mm*GMAS with N-terminal 6xHis-Tag was cloned in the pET28a(+) plasmid, by NcoI and NotI restriction digestion and following ligation. Recombinant *Mm*GMAS was produced in *E. coli* BL21 (DE3) (New England Biolab, United States) transformed with pET28_*Mm*GMAS. Preparative cultures were carried out in 1 L of LB medium, and the cells were grown in a shaking incubator (180 rpm) at 37 °C to an optical density at 600 nm (OD_600_) of 0.4–0.6. Protein expression was induced by adding 0.2 mM isopropyl-β-D-1-thiogalactopyranoside (IPTG) and prolonged overnight at room temperature. Cells were harvested by centrifugation (4 °C, 15 min, 5000×g) and resuspended in 50 mM sodium phosphate pH 7.0, 10% v/v glycerol. Cell lysis was obtained by a French Press (One Shot Cell Disruptor; Constant Systems, United Kingdom). The crude extract was centrifuged at 4 °C for 30 min at 40000×g before loading the soluble fraction on a HisTrap^TM^ HP 1 mL column (Cytiva, United States), pre-equilibrated in 50 mM sodium phosphate pH 7.0, 10% v/v glycerol. Following extensive washes in the same buffer, 6xHis-*Mm*GMAS was eluted by 50 mM sodium phosphate pH 7.0, 300 mM imidazole, 10% v/v glycerol (0%-100% linear gradient in 30 column volumes) using a GE AKTA Purifier 100 FPLC System. Fractions containing mostly pure *Mm*GMAS, as revealed by SDS-PAGE analysis (Fig. S12), were pooled and buffer-exchanged to 50 mM sodium phosphate pH 7.0, 10% v/v glycerol using PD-10 desalting columns (Cytiva, United States). The 6xHisTag was removed by thrombin cleavage, incubating 6xHis-*Mm*GMAS overnight with thrombin (Merck Life Sciences, Germany), 20:1 weight ratio. Then, *Mm*GMAS was further purified by Size Exclusion Chromatography (SEC) using a Superdex 200 26/60 column (Cytiva, United States) in the same buffer.

### Differential Scanning Fluorimetry (DSF)

Differential Scanning Fluorimetry (DSF) was performed to evaluate the apparent unfolding temperature of recombinant *Mm*GMAS in different buffers (50 mM Tris pH 8.0, PBS pH 7.0 and Tricine pH 7.5) and in the presence of different additives (15% and 30% v/v glycerol and ethylene glycol; 5% and 10% w/v sucrose). Measurements were performed in duplicate using the exogenous SYPRO orange fluorogenic dye (Thermo Fisher Scientific, United States) and according to the protocol previously described in Fogal et al., 2015.

### Protein crystallization and X-ray data analysis

Purified *Mm*GMAS was concentrated to 12 mg/mL by a Vivaspin (Sartorius, United Kingdom) ultrafiltration centrifugal device and supplemented by 3 mM MgCl_2_ and 1.4 mM ATPγS, before dispensing large protein crystallization screenings (PACT Premier, Morpheus, JCSG Plus, LMB; Molecular Dimensions Ltd., United Kingdom) in the sitting-drop isothermal vapor diffusion setup. Drops containing equal volumes of protein and reservoir solutions (0.4 µL total volume) were dispensed on MRC 96-Well 2-Drop Crystallization plates, and incubated at 20 °C. The best-diffracting crystals were obtained in the Morpheus A2 precipitant buffer (0.03 M MgCl_2_⋅6H_2_O, 0.03 M CaCl_2_⋅2H_2_O, 20% v/v Ethylene glycol, 10 % w/v PEG 8000, 0.1 M Imidazole, 0.1 M MES monohydrate, pH 6.5) and were not further cryoprotected before freezing in liquid nitrogen. X-ray diffraction data (doi: 10.15151/ESRF-ES-1581819979) were collected at the ID30A-3 beamline of the European Synchrotron Radiation Facility (ESRF, Grenoble, France). After several trials, a dataset in the C 1 2 1 space group and automatically processed by the Xia2/DIALS pipeline (Winter et al., 2022) was phased by molecular replacement via Molrep (Vagin & Teplyakov, 2010) using an AlphaFold-generated model of *Mm*GMAS as the starting model (Jumper et al., 2021).

Further data reduction was carried out by Aimless (Evans, 2011) via the CCP4i2 interface (Agirre et al., 2023). The molecular model was refined automatically by PDB-Redo (Joosten et al., 2014) and REFMAC5 (Murshudov et al., 2011), and manually by Coot (Emsley et al., 2010).

6 Protein chains (A-F) are present in the asymmetric unit (ASU) and clearly visible in the electron density map from residue Ser1 (chains A-C), Glu2 (chain D) or Met1 (chains E, F), to the C-terminal residue Tyr444. The homo-dodecameric biological assembly of *Mm*GMAS can be reconstructed by applying the crystallographic symmetry operators. While chains A-D of the ASU belong to the same dodecamer, chains E and F belong to an adjacent one.

Protein-protein and protein-ligand interactions were analyzed by PDBePISA (https://www.ebi.ac.uk/pdbe/pisa/) (Krissinel & Henrick, 2007) and PLIP (https://plip-tool.biotec.tu-dresden.de/plip-web/plip/index) (Adasme et al., 2021). Surface electrostatic potential was calculated by APBS-PDB2PQR (https://server.poissonboltzmann.org/) (Jurrus et al., 2018), simulating an environment with pH 7.0 and 0.15 M NaCl.

### Modelling and molecular dynamics simulations

2 *Mm*GMAS subunits belonging to the same hexameric ring were modeled in complex with 3 Mg^2+^, ADP, methylamine (MMA) and L-γ-glutamyl phosphate (P-γ-GLU) by Boltz-1 (Wohlwend et al., 2024) via the Tamarind Bio web portal (https://www.tamarind.bio/). To ensure that a dimer of laterally interacting *Mm*GMAS chains was modelled and to reduce the computational resources for subsequent molecular dynamics, *Mm*GMAS sequence was trimmed at residue 424.

Conversely, to model MMA in the previously proposed binding site (Wang et al., 2021b), it was docked on *Mm*GMAS-ADP-P-γ-GLU by AutoDock Vina (Bugnon et al., 2024) via the SwissDock webserver (Eberhardt et al., 2021).

Molecular dynamics simulations were performed by Gromacs 2022.3 (Berendsen et al., 1995; Abraham et al., 2015) using the Charmm36-jul2021 forcefield (Best et al., 2012). Ligands (MMA, ADP and P-γ-GLU) were parametrized by CGenFF (Vanommeslaeghe et al., 2010). Simulations of 800 ns with explicit solvent were performed as previously described (Chinellato et al., 2023; Lemkul, 2018). Briefly, the models were solvated in a rectangular box using the TIP3P water model, ensuring a minimum distance of 1 nm between the protein complex and the box boundaries. 0.15 M Na^+^ and Cl^-^ ions were added to neutralize the system net charge and simulate a physiological ionic strength. The system was first minimized with a tolerance of 1000 kJ mol^−1^ nm^−2^ allowing a maximum of 500000 iterations of steepest descent. Subsequently, the system was heated from 0 to 100 K during a 200 ps NVT MD simulation with positional restraints of 400 kJ mol^−1^ nm^−2^. Then, the system was heated up to 310 K in 400 ps during an NPT simulation with further lowered restraint (200 kJ mol^−1^ nm^−2^) and further equilibrated over a 5 ns NPT simulation with backbone restraints of 50 kJ mol^−1^ nm^−2^. All restraints were removed for the 800 ns production run. The V-rescale thermostat was used to equilibrate the temperature, whereas the C-rescale barostat was used to control the pressure. Newton’s equation of motion was integrated using a leapfrog algorithm with a 2 fs time step. The particle mesh Ewald (PME) method was used to compute the long-range electrostatic force.

Rotational and translational motions of the system were removed, and all bonds were constrained with the LINCS algorithm.

Simulation convergence was assessed by analyzing the global Root Mean Square Deviation (RMSD). Interatomic distances were calculated using the Gromacs built-in pairdist tool.

## AKNOWLEDGMENTS

The work was supported by fundings from the Department of Biology, University of Padova (projects BIRD221528/22 and U-GOV CEND_BIRD24_01), that also provided the equipment of the Plant Genome Editing and Phenotyping Facility for cyanobacteria cultivations.

The authors would like to thank the staff of beamline ID30A-3 of the European Synchrotron Radiation Facility (ESRF, Grenoble, France) for technical assistance during X-ray diffraction data collection. Moreover, Daniele de Sanctis (ESRF, Grenoble, France) is acknowledged for insightful discussions.

## REFERENCES

1. Abraham, M. J. et al. Gromacs: High performance molecular simulations through multi-level parallelism from laptops to supercomputers. SoftwareX 1–2, 19–25 (2015).

2. Adasme, M. F. et al. PLIP 2021: Expanding the scope of the protein-ligand interaction profiler to DNA and RNA. Nucleic Acids Research 49, W530–W534 (2021). Abraham, M. J. et al. Gromacs: High performance molecular simulations through multi-level parallelism from laptops to supercomputers. *SoftwareX* 1–2, 19–25 (2015).

3. Agirre, J. et al. The CCP4 suite: integrative software for macromolecular crystallography. Acta Crystallographica Section D: Structural Biology 79, 449–461 (2023).

4. Andexer, J. N. & Richter, M. Emerging enzymes for ATP regeneration in biocatalytic processes. ChemBioChem 16, 380–386 (2015).

5. Benninghaus, L., Walter, T., Mindt, M., Risse, J. M. & Wendisch, V. F. Metabolic Engineering of *Pseudomonas putida* for Fermentative Production of l-Theanine. Journal of Agricultural and Food Chemistry 69, 9849–9858 (2021).

6. Berendsen, H. J. C., van der Spoel, D. & van Drunen, R. GROMACS: A message-passing parallel molecular dynamics implementation. Comput. Phys. Commun. 91, 43–56 (1995).

7. Best, R. B. et al. Optimization of the additive CHARMM all-atom protein force field targeting improved sampling of the backbone φ, ψ and side-chain χ1 and χ2 Dihedral Angles. J. Chem. Theory Comput. 8, 3257–3273 (2012).

8. Bugnon, M. et al. SwissDock 2024: major enhancements for small-molecule docking with Attracting Cavities and AutoDock Vina. Nucleic Acids Res. 52, W324–W332 (2024).

9. Chen, Y. et al. Effects of long-term nitrogen fertilization on the formation of metabolites related to tea quality in subtropical China. Metabolites 11, 1–22 (2021).

10. Chinellato, M. et al. 1-Piperidine Propionic Acid as an Allosteric Inhibitor of Protease Activated Receptor-2. Pharmaceuticals 16, 1–14 (2023).

11. Dossena, L., Curti, B. & Vanoni, M. A. Activation and coupling of the glutaminase and synthase reaction of glutamate synthase is mediated by E1013 of the ferredoxin-dependent enzyme, belonging to loop 4 of the synthase domain. Biochemistry 46, 4473–4485 (2007).

12. Eberhardt, J., Santos-Martins, D., Tillack, A. F. & Forli, S. AutoDock Vina 1.2.0: New Docking Methods, Expanded Force Field, and Python Bindings. J. Chem. Inf. Model. 61, 3891–3898 (2021).

13. Ekanayake, A; Li, J. J. (12) United States Patent. US7303773, 1–8 (2007).

14. Emsley, P., Lohkamp, B., Scott, W. G. & Cowtan, K. Features and development of Coot. Acta Crystallographica Section D: Biological Crystallography 66, 486–501 (2010).

15. Evans, P. R. An introduction to data reduction: Space-group determination, scaling and intensity statistics. Acta Crystallographica Section D: Biological Crystallography 67, 282–292 (2011).

16. Fan, X. et al. Pathway engineering of *Escherichia coli* for one-step fermentative production of L-theanine from sugars and ethylamine. Metabolic Engineering Communications 11, e00151 (2020).

17. Fan, C., Qi, J., Cong, Y. & Zhang, C. Enhanced L-theanine production through semi-rational design of γ-glutamylmethylamide synthetase from Methylovorus mays. Enzyme Microb. Technol. 180, 110481 (2024).

18. Fogal, S., Beneventi, E., Cendron, L. & Bergantino, E. Structural basis for double cofactor specificity in a new formate dehydrogenase from the acidobacterium *Granulicella mallensis* MP5ACTX8. Applied Microbiology and Biotechnology 99, 9541–9554 (2015).

19. Fu, X. et al. Characterization of l-theanine hydrolase in vitro and subcellular distribution of its specific product ethylamine in tea (*Camellia sinensis*). Journal of Agricultural and Food Chemistry 68, 10842–10851 (2020).

20. Hidese, S., Ogawa, S., Ota, M., Ishida, I. & Yasukawa, Z. Effects of L-Theanine Administration on Stress-Related Symptoms and Cognitive Functions in. Nutrients 11, 1–13 (2019).

21. Joosten, R. P., Long, F., Murshudov, G. N. & Perrakis, A. The PDB-REDO server for macromolecular structure model optimization. IUCrJ 1, 213–220 (2014).

22. Jumper, J. et al. Highly accurate protein structure prediction with AlphaFold. Nature 596, 583–589 (2021).

23. Jurrus, E. et al. Improvements to the APBS biomolecular solvation software suite. Protein Science 27, 112–128 (2018).

24. Kimura, K., Ozeki, M., Juneja, L. R. & Ohira, H. l-Theanine reduces psychological and physiological stress responses. Biological Psychology 74, 39–45 (2007).

25. Klughammer, C. & Schreiber, U. Saturation Pulse method for assessment of energy conversion in PS I. PAM Application Notes 11–14 (2008).

26. Krissinel, E. & Henrick, K. Inference of Macromolecular Assemblies from Crystalline State. Journal of Molecular Biology 372, 774–797 (2007).

27. Lemkul, J. From Proteins to Perturbed Hamiltonians: A Suite of Tutorials for the GROMACS-2018 Molecular Simulation Package. Living J. Comput. Mol. Sci. 1, 1–53 (2019).

28. Li, Y., Horsman, M., Wu, N., Lan, C. Q. & Dubois-Calero, N. Biofuels from microalgae. In Biotechnology Progress vol. 24 815–820 (2008).

29. Li, F. et al. Theanine transporters are involved in nitrogen deficiency response in tea plant (*Camellia sinensis* L.). Plant Signaling and Behavior 15, 20–23 (2020).

30. Li, H. & Sherman, L. A. Characterization of *Synechocystis* sp. strain PCC 6803 and Δnbl mutants under nitrogen-deficient conditions. Archives of Microbiology 178, 256–266 (2002).

31. Loprete, G., et al. Greening the production of indigo blue exploiting light and a recombinant *Synechocystis* sp. PCC6803 strain expressing the enzyme mFMO. Microb. Biotechnol. 18, e70146 at 10.1111/1751-7915.70146 (2025).

32. Ma, H. et al. Efficient fermentative production of l-theanine by *Corynebacterium glutamicum*. Applied Microbiology and Biotechnology 104, 119–130 (2020).

33. Malihan-Yap, L., Grimm, H. C. & Kourist, R. Recent Advances in Cyanobacterial Biotransformations. Chemie-Ingenieur-Technik vol. 94, 1628–1644 at 10.1002/cite.202200077 (2022).

34. Matsumoto, K. et al. Antidiabetic activity of Zn(II) complexes with a derivative of L-glutamine. Bulletin of the Chemical Society of Japan 78, 1077–1081 (2005).

35. Murshudov, G. N. et al. REFMAC5 for the refinement of macromolecular crystal structures. Acta Crystallographica Section D: Biological Crystallography 67, 355–367 (2011).

36. Noreña-Caro, D. & Benton, M. G. Cyanobacteria as photoautotrophic biofactories of high-value chemicals. Journal of CO2 Utilization vol. 28 335–366 at 10.1016/j.jcou.2018.10.008 (2018).

37. Pan, X. et al. Efficient synthesis of γ-glutamyl compounds by co-expression of γ-glutamylmethylamide synthetase and polyphosphate kinase in engineered *Escherichia coli*. Journal of Industrial Microbiology and Biotechnology 47, 573–583 (2020).

38. Park, J. & Choi, Y. Cofactor engineering in cyanobacteria to overcome imbalance between NADPH and NADH: A mini review. Frontiers of Chemical Science and Engineering 11, 66–71 (2017).

39. Patil, P. P., Vass, I., Kodru, S. & Szabó, M. A multi-parametric screening platform for photosynthetic trait characterization of microalgae and cyanobacteria under inorganic carbon limitation. PLoS ONE 15, 1–25 (2020).

40. Perucho, J. et al. Optimal excitation and emission wavelengths to analyze amino acids and optimize neurotransmitters quantification using precolumn OPA-derivatization by HPLC. Amino Acids 47, 963–973 (2015).

41. Pope, M. A., Hodge, J. A. & Nixon, P. J. An Improved Natural Transformation Protocol for the Cyanobacterium *Synechocystis* sp. PCC 6803. Front. Plant Sci. 11, (2020).

42. Ravasio, S. et al. Properties of the recombinant ferredoxin-dependent glutamate synthase of *Synechocystis* PCC6803. Comparison with the *Azospirillum brasilense* NADPH-dependent enzyme and its isolated α subunit. Biochemistry 41, 8120–8133 (2002).

43. Rosgaard, L., de Porcellinis, A. J., Jacobsen, J. H., Frigaard, N. U. & Sakuragi, Y. Bioengineering of carbon fixation, biofuels, and biochemicals in cyanobacteria and plants. Journal of Biotechnology 162, 134–147 (2012).

44. Schmelling, N. M. & Bross, M. What is holding back cyanobacterial research and applications? A survey of the cyanobacterial research community. Nature Communications 15, 1–9 (2024).

45. Sharma, E., Joshi, R. & Gulati, A. L-Theanine: An astounding sui generis integrant in tea. Food Chemistry 242, 601–610 (2018).

46. Tavanti, M., Hosford, J., Lloyd, R. C. & Brown, M. J. B. Recent Developments and Challenges for the Industrial Implementation of Polyphosphate Kinases. ChemCatChem 13, 3565–3580 (2021).

47. Tsushida, T. & Takeo, T. An enzyme hydrolyzing L-theanine in tea leaves. Agricultural and Biological Chemistry 49, 2913–2917 (1985).

48. Vagin, A. & Teplyakov, A. Molecular replacement with MOLREP. Acta Crystallographica Section D: Biological Crystallography 66, 22–25 (2010).

49. Vanommeslaeghe, K. et al. CHARMM general force field: A force field for drug-like molecules compatible with the CHARMM all-atom additive biological force fields. Journal of computational chemistry, 31(4): 671–690 (2010)

50. Wang, Q. et al. L-theanine as a promising agent on brain health-promoting foods – A review. Journal of Food Bioactives 13, 32–39 (2021a).

51. Wang, N. et al. Crystal structures of-glutamylmethylamide synthetase provide insight into bacterial metabolism of oceanic monomethylamine. Journal of Biological Chemistry 296, 100081 (2021b).

52. Waterbury, J. B., Watson, S. W., Guillard, R. R. L., & Brand, L. E. Widespread occurrence of a unicellular, marine, planktonic, cyanobacterium. Nature 277, 293–294 (1979).

53. Winter, G. et al. DIALS as a toolkit. Protein Science 31, 232–250 (2022).

54. Wohlwend, J. et al. Boltz-1: Democratizing Biomolecular Interaction Modeling. bioRxiv, 2024-11 at 10.1101/2024.11.19.624167 (2024).

55. Yamada, T. et al. Administration of theanine, a unique amino acid in tea leaves, changed feeding-relating components in serum and feeding behavior in rats. *Bioscience*, Biotechnology and Biochemistry 72, 1352–1355 (2008).

56. Yamamoto, S., Wakayama, M. & Tachiki, T. Cloning and expression of *Methylovorus mays* no. 9 gene encoding γ-glutamylmethylamide synthetase: An enzyme usable in theanine formation by coupling with the alcoholic fermentation system of baker’s yeast. Bioscience, Biotechnology and Biochemistry 72, 101–109 (2008).

57. Yang, T. et al. Combining protein and metabolic engineering strategies for high-level production of L-theanine in *Corynebacterium glutamicum*. Bioresource Technology 394, 130200 (2024).

58. Yoto, A., Motoki, M., Murao, S. & Yokogoshi, H. Effects of L-theanine or caffeine intake on changes in blood pressure under physical and psychological stresses. Journal of physiological anthropology 31, 28 (2012).

59. Zhou, J., Zhang, F., Meng, H., Zhang, Y. & Li, Y. Introducing extra NADPH consumption ability significantly increases the photosynthetic efficiency and biomass production of cyanobacteria. Metabolic Engineering 38, 217–227 (2016).

60. Zhou, J., Zhou, J. X., Yang, H. M., Yan, C. S. & Huang, F. Characterization of a sodium-regulated glutaminase from cyanobacterium *Synechocystis* sp. PCC 6803. Sci. China, Ser. C Life Sci. 51, 1066–1075 (2008).

