## Supporting Information for "Harnessing photosynthetic ATP for whole-cell biocatalysis in the cyanobacterium *Synechocystis*"

**Supplementary Results**

**Construction of the transgenic strain *Syn*_GMAS**

The sequence coding for γ-Glutamyl-MethylAmide Synthetase from the methylotrophic bacteria *Methylovorus mays* No. 9 (*Mm*GMAS, accession number: A9ZPH9) was designed to permit its expression in fusion with a N-terminal 6XHis-Tag, and with NcoI and NotI restriction sites at the 5' and 3' termini respectively, useful for cloning it into the integrative vector pSuperP_UV for expression in *Synechocystis* (Loprete et al., 2025). The resulting plasmid, pSuperP_*Mm*GMAS, was used for the construction of the homoplasmic and stable *Synechocystis* strain named *Syn*_GMAS. In its chromosome, the expression of the enzyme is driven by the well characterized "SuperPromoter" Pcpc560, a native *Synechocystis* promoter frequently employed for producing recombinant proteins in the cyanobacterium. Together with the kanamycin resistance cassette also present in the vector (Fig. S1), the construct interrupts the locus ssl0410, which is a known neutral site of the cyanobacterial chromosome. PCR confirmed correct insertion and achievement of homoplasmy (Fig. S2); the analysis was performed by comparing amplimers from the genomic DNA of *Syn*_GMAS with those obtained from the genomes of both wild-type and *Syn*_UV strains, the latter one being a "pseudo" wild-type strain containing the pSuperP_UV "empty" construct, only carrying the antibiotic resistance cassette (Loprete et al., 2025). Production of the protein was verified by Western blotting (Fig. S3).

**In vitro synthesis of L-theanine catalyzed by the recombinant *Mm*GMAS produced in *E. coli***

Sodium glutamate (50 mM, 18.7 mg), ethylamine hydrochloride (50 mM, 8.2 mg), $MgCl_2$ (50 mM, 9.5 mg) and ATP (50 mM, 55.1 mg) were solubilized in 100 mM Trizma buffer spontaneous pH (1.5 mL). The pH was adjusted to 7.5 with NaOH (10 M). The reaction was initiated by addition of *Mm*GMAS (500 µL, 1 mg/mL) and incubated at 37 °C under vigorous mixing (1000 rpm). During the reaction the pH was checked by pH indicator paper and if necessary adjusted to 7.5 with NaOH (10 M). At different endpoints, samples were withdrawn (20 µL) from the reaction mixture, filtered with Microcon® filter (cutoff 10 kDa) for 5 min at 13,000 rpm and 25 °C to eliminate the enzyme, derivatized with OPA reagent by following the standard protocol reported in the Methods section and analyzed by HPLC. Conversion was determined by using a calibration curve of an authentic sample of L-thea. Conversion of L-thea is equal to 84% (Fig. S6).

**L-theanine stability under reaction condition in the absence or presence of recombinant *Mm*GMAS**

L-theanine (50 mM, 17.4 mg), $MgCl_2$ (50 mM, 9.5 mg) and ATP (50 mM, 55.1 mg) were solubilized in 100 mM Trizma buffer spontaneous pH (2 mL). The pH was adjusted to 7.5 with NaOH (10 M). Following the addition of *Mm*GMAS (500 µL, 1 mg/mL) or buffer (500 µL), samples were incubated at 37 °C under vigorous mixing (1000 rpm). At different endpoints, samples were withdrawn (20 µL) from the reaction mixture, derivatized with OPA reagent by following the standard protocol reported in the Methods section and analyzed by HPLC. L-thea is stable under the tested reaction conditions, both in the absence of the recombinant enzyme (Fig. S8A) and when it is added to the mixture (Fig. S8B).

**Details of structural characterization, docking and molecular dynamics simulations of *Mm*GMAS**

*Mm*GMAS is organized as a homo-dodecamer composed of 2 stacked hexameric rings. Adjacent chains keep the two rings together by swapping the C-terminal α-helix, which docks on a wide cleft on the partner monomer (Fig. S13B). This interaction hides on average 2358 $Å^2$ on each protein chain, while an average surface of 1393 $Å^2$ is buried between neighboring chains in the same hexameric ring (Fig. S13A). Both kinds of interaction involve several electrostatic and hydrophobic contacts. One ATPγS molecule is coordinated by each *Mm*GMAS monomer, as clearly defined in the electron density map (Fig. 5B). Three $Mg^{2+}$ cations have been assigned to electron density peaks near the ATPγS, and their position validated by the well-known role of this ion in the stabilization of nucleotide cofactors, and by comparison with previously solved structures of *Rh*GMAS (PDB: 7CQU, 7CQQ; Wang et al., 2021). ATPγS in *Mm*GMAS assumes a conformation similar to that of AMPPNP in complex with *Rh*GMAS (PDB 7CQQ) and the residues lining the ATP-binding site are conserved among the two orthologues. In particular, the adenine base of ATPγS forms π-stacking and π-cation interactions with Tyr198 and Arg331 sidechains, respectively. ATPγS is further stabilized by H-bonds with Tyr181, Asn195, Tyr196, Tyr198, Ser246, Arg331 and Arg335. The negatively charged phosphate groups are ion-coupled to His244, Arg321 and Arg326 and coordinate the $Mg^{2+}$ cations along with Glu126, Glu128, Glu186, Glu193, His242 and Glu333. The octahedral coordination sphere of $Mg^{2+}$ cations is completed by water molecules, which are not visible in all the chains due to local variability and averaging effects.

Superimposition of the 6 *Mm*GMAS monomers in the ASU demonstrated a high conformational homogeneity (RMSD < 0.07 Å), except for the region Leu298-Leu304, which explores slightly different conformations, coherent with the observed higher B factors. These residues belong to a flexible loop, which was described to undergo a conformational change upon ATP binding (Wang et al., 2021).

Each *Mm*GMAS chain defines a catalytic cleft with the contribution of the adjacent chain within the same hexameric ring. This cleft comprises a broad, slightly positively charged region for adenine nucleotide binding, three small negatively charged cavities, and a deeper, highly positively charged pocket which was shown to accommodate the glutamate substrate (Fig. 5C; Wang et al., 2021). Based on electrostatic complementarity, Wang and colleagues proposed that one of the lateral chambers in the *Rh*GMAS cleft binds methylamine, with Asp177 and Glu186 acting as catalytic bases. This was supported by mutagenesis of putative binding residues, though no structural or computational evidence directly confirmed MMA binding site.

Then, to gain more insights on substrates binding by *Mm*GMAS, we modelled it in the quaternary complex including ADP, its natural substrate monomethylamine (MMA) and ʟ-γ-glutamyl phosphate (P-γ-Glu; Fig. 5C-D), since the catalytic mechanism implies that MMA binds GMAS after the transfer of the γ-phosphate group from ATP to ʟ-glutamate. In the generated model, ADP and P-γ-Glu were observed closely aligned with experimentally solved adenine cofactors and substrate mimics in both *Mm*GMAS (PDB 9QUR) and *Rh*GMAS (PDB 7CQU). Notably, the ionizable groups of P-γ-Glu are stabilized by electrostatic interactions with Arg321, Arg335, Arg297, His242 and Glu128. While the *Mm*GMAS model aligns well with our crystallographic structure at both global and local levels, the presence of the amine substrate induces a slight conformational shift in Asp187, positioning it favorably for coordination with MMA. The spatial localization of *Mm*GMAS Asp187 and Glu186 (Asp180 and Glu179 in *Rh*GMAS) with respect to MMA and P-γ-Glu suggests that they can be the catalytic bases needed for MMA and tetrahedral intermediate deprotonation. However, in our analysis, methylamine (MMA) was observed to occupy a slightly different position than previously hypothesized by Wang and collaborators, notably closer to the Cδ of P-γ-Glu, and properly positioned to serve as a nucleophile (Wang et al., 2021). In the model we propose, MMA is stabilized through hydrophobic interactions with residues Ala51′ and Ala54′ of the adjacent *Mm*GMAS chain, complemented by electrostatic contacts between MMA amine group (protonated at physiological pH) and the acidic side chains of Glu186 and Asp187 (Fig. 5D). To further validate this binding hypothesis, we conducted molecular dynamics simulations to compare the stability of MMA both in the proposed binding site and the *Rh*GMAS model. Notably, MMA remained stable in our predicted complex for a higher and adequate time frame for exerting the nucleophilic attack on P-γ-Glu (170 ns; Fig. S14). In contrast, MMA docked in the previously proposed binding pocket remained stable for a significantly shorter duration (20 ns). These computational observations suggest that our novel described binding mode for MMA may be crucial for GMAS catalysis.

Our hypothesis aligns with previous site-directed mutagenesis studies on *Rh*GMAS that targeted residues potentially involved in substrate binding and catalysis. Notably, mutation of Glu179 (Wang et al., 2021), corresponding to Glu186 in *Mm*GMAS, was found to abolish catalytic activity. This underscores the critical role of Glu186 and Asp187 in enabling the proper positioning of the amine substrate for effective catalysis and/or their catalytic role in MMA and tetrahedral intermediate deprotonation. The convergence of these findings highlights the significance of these residues in optimizing the catalytic process and reinforces the relevance of our proposed model.

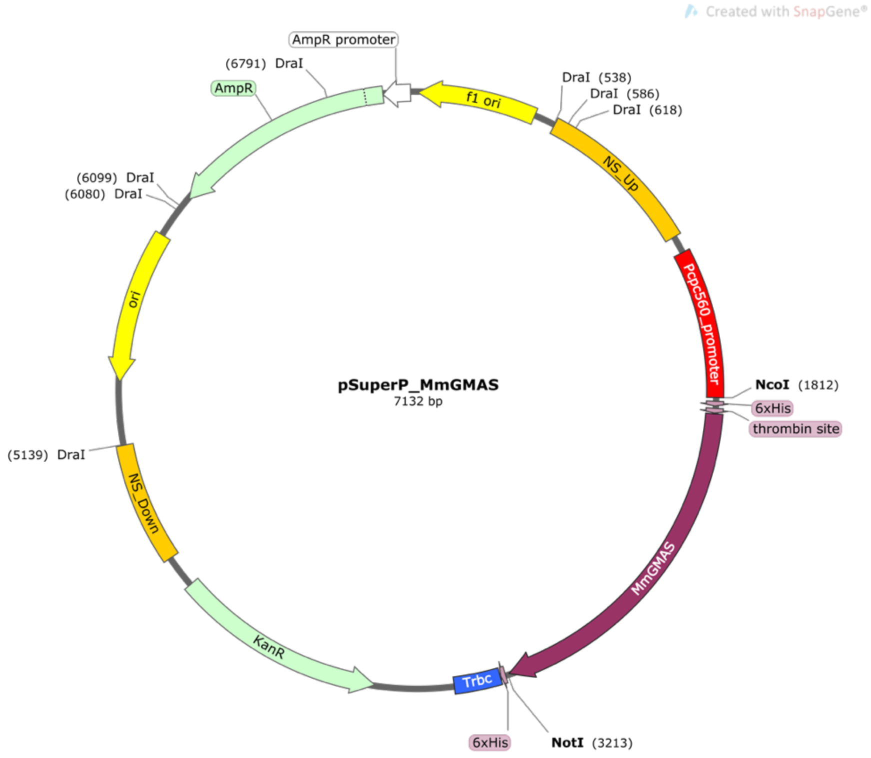

**Fig. S1. Map of the plasmid pSuperP_*Mm*GMAS.** Map obtained by the Snapgene® software (Dotmatics; available at snapgene.com), included with the NcoI and NotI restrictrion sites used for *Mm*GMAS coding sequence cloning and DraI restriction sites used for DNA linearization prior to *Synechocystis* transformation

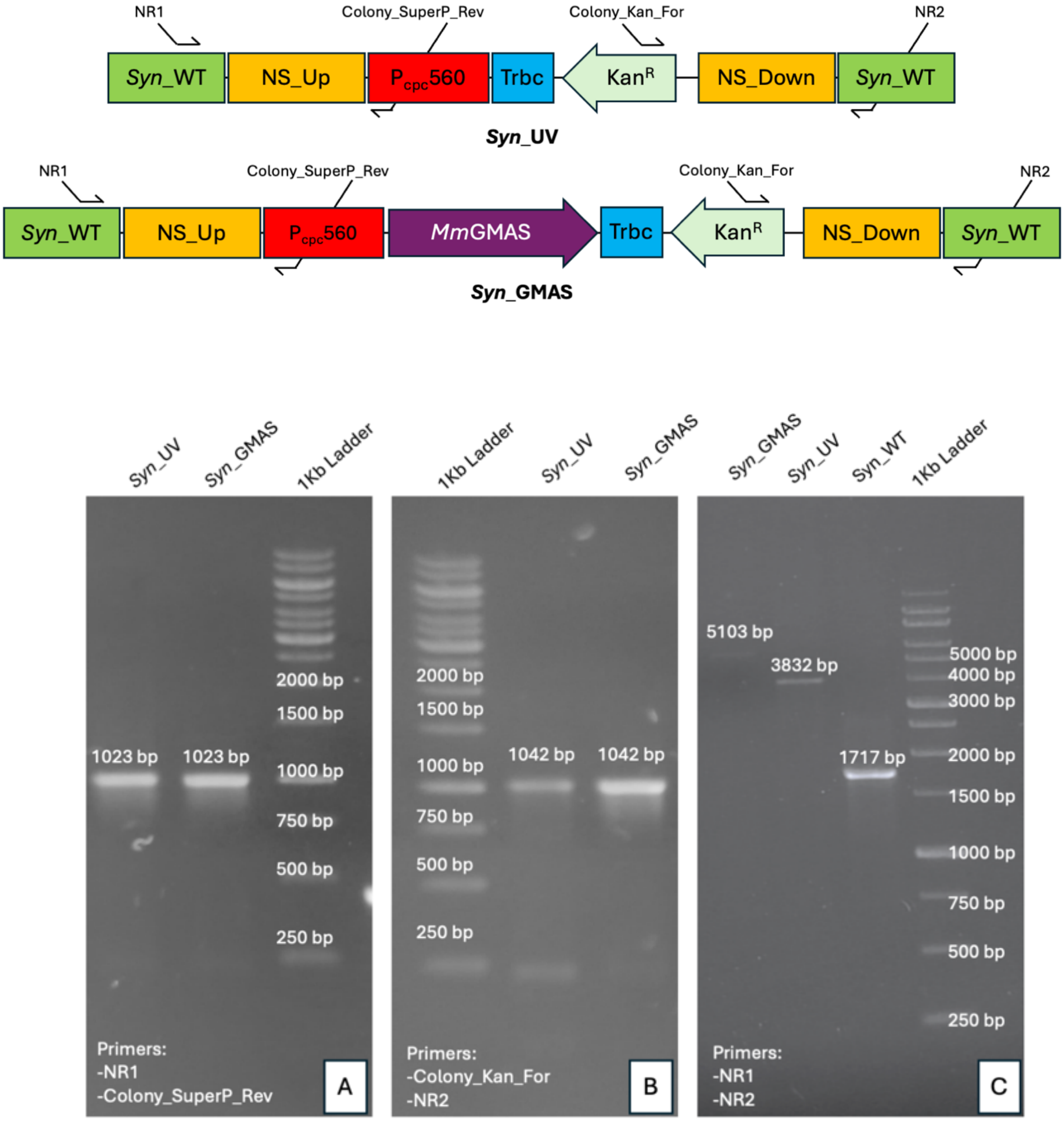

**Fig. S2. PCR amplification of the transgenic strain *Syn*_GMAS chromosome**. Agarose gel electrophoresis image of colony PCR amplifications used for verifying (i) correct insertion of the engineered sequence (A and B) and (ii) achievement of homoplasmy in both *Syn*_UV and *Syn*_GMAS strains (C). The diagram above the agarose gel image illustrates the map of the chromosomal region of interest, as expected after transformation and recombination. The locations of the primers used for amplification are indicated, with green segments representing the cyanobacterial chromosome regions flanking the selected neutral insertion site (*Syn*_WT). The scheme does not respect the base pair lengths.

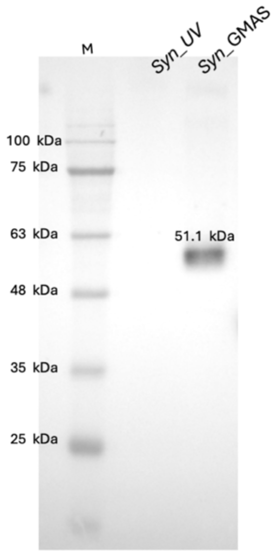

**Fig. S3. Western-Blot analysis of *Syn*_GMAS and Syn_UV protein extracts.** Total protein extracts were obtained from cells cultivated in standard BG11 growth medium and conditions. Immunodetection was performed by using an anti-His-Tag-HRP antibody (SB194b, Southern Bio, Birmingham, USA). The indicated molecular weight of *Mm*GMAS is calculated from the primary sequence of the His-tagged protein.

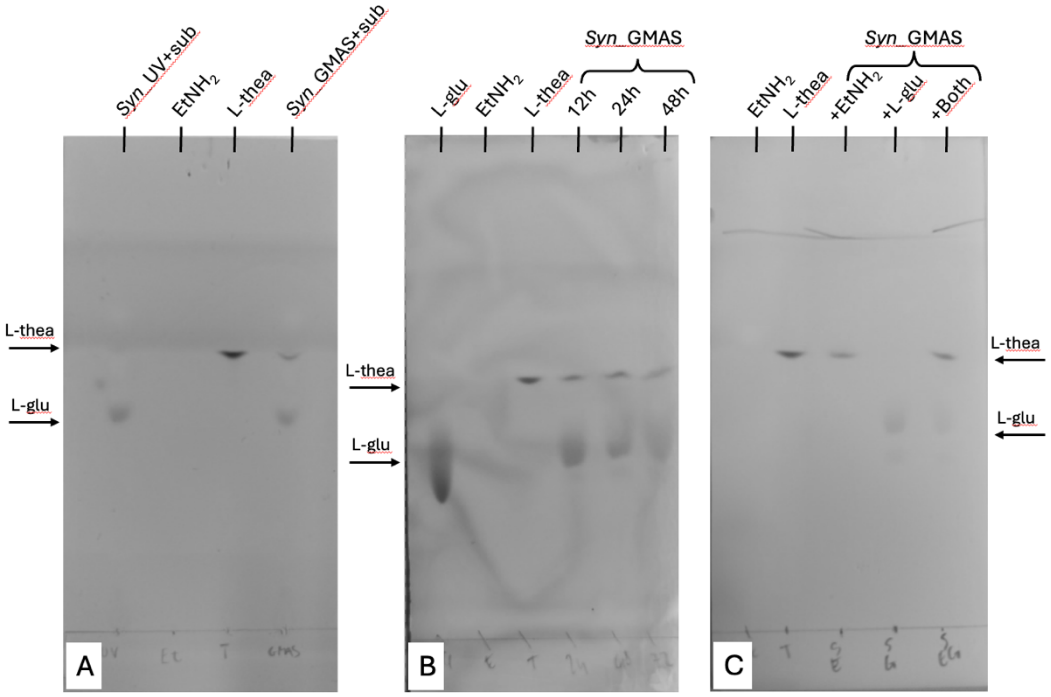

**Fig. S4. TLC analysis of whole-cell biotransformations.** Mobile phase used for separation was composed of 70/30 $NH_3$/EtOH, and detection was performed by 0.1% w/v ninhydrin in EtOH followed by 2 minutes at 60 °C. Biotransformations were performed in the presence of 1 mM of each substrate.

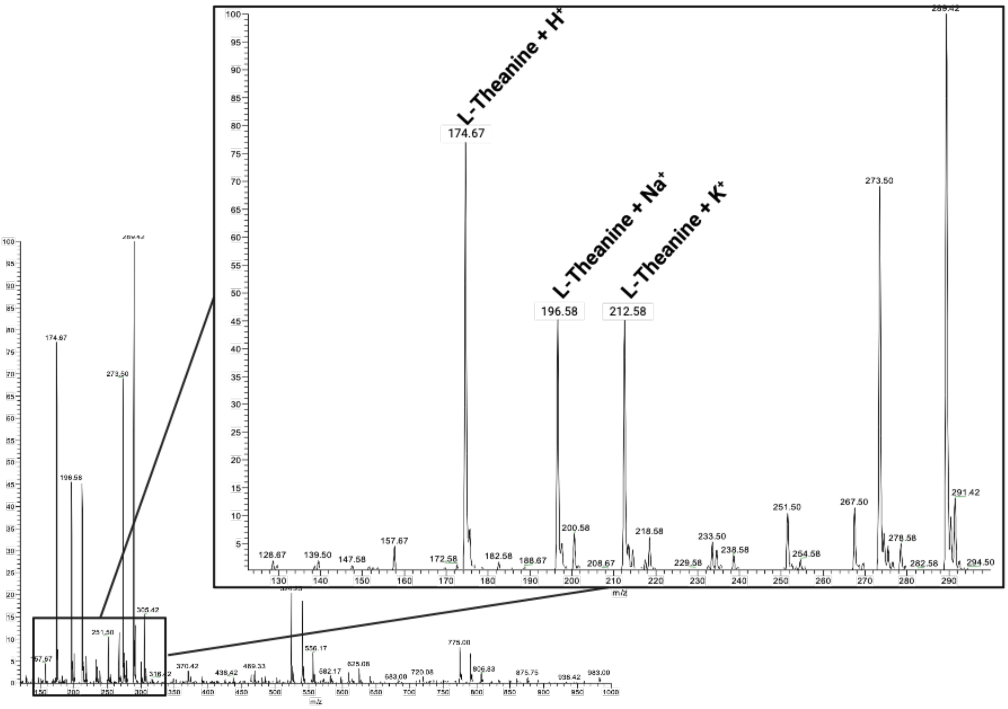

**Fig. S5. Mass Spectrometry (MS) analysis of biotransformation supernatant**. Analysis was performed on the supernatant of a 48 hours whole-cell biotransformation by *Syn*_GMAS supplemented with 1 mM L-glutamate and ethylamine.

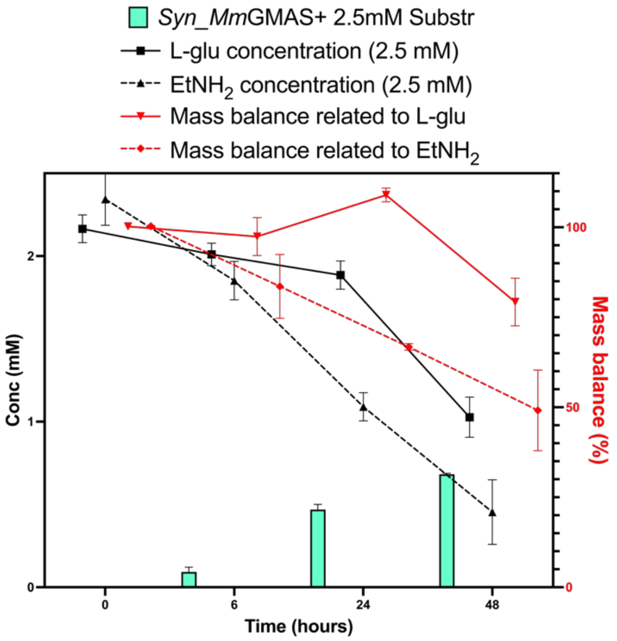

**Fig. S6. Substrates concentration and mass balance of *Syn*_GMAS whole-cell biotransformations.** Results of RP-HPLC analysis of supernatant aliquots from *Syn*_GMAS whole-cell biotransformations at different endpoints, supplemented with 2.5 mM L-glutamate and ethylamine. Black lines represent evolution of L-glutamate and ethylamine over time. Red lines indicate the mass balance, calculated as the separate sums of L-theanine and L-glutamate or ethylamine. Biotransformation and analysis were performed in triplicate.

| time (h) | Area Glu | Area Ser | Area Thea | area $EtNH_2$ | Conv. (%) |
|---|---|---|---|---|---|
| 0 | 2093,932 | 205,119 | 0 | 1985,596 | 0 |
| 2 | 1450,634 | 206,245 | 787,951 | 1319,82 | 35.92 |
| 4 | 1068,773 | 203,071 | 1211,002 | 883,886 | 55.65 |
| 6 | 988,938 | 203,852 | 1577,244 | 826,12 | 72.73 |
| 8 | 890,384 | 214,222 | 1688,282 | 697,88 | 77.91 |
| 24 | 492,82 | 205,643 | 1800,646 | 310,577 | 83.15 |
| 48 | 368,628 | 209,069 | 1815,307 | 211,035 | 83.83 |

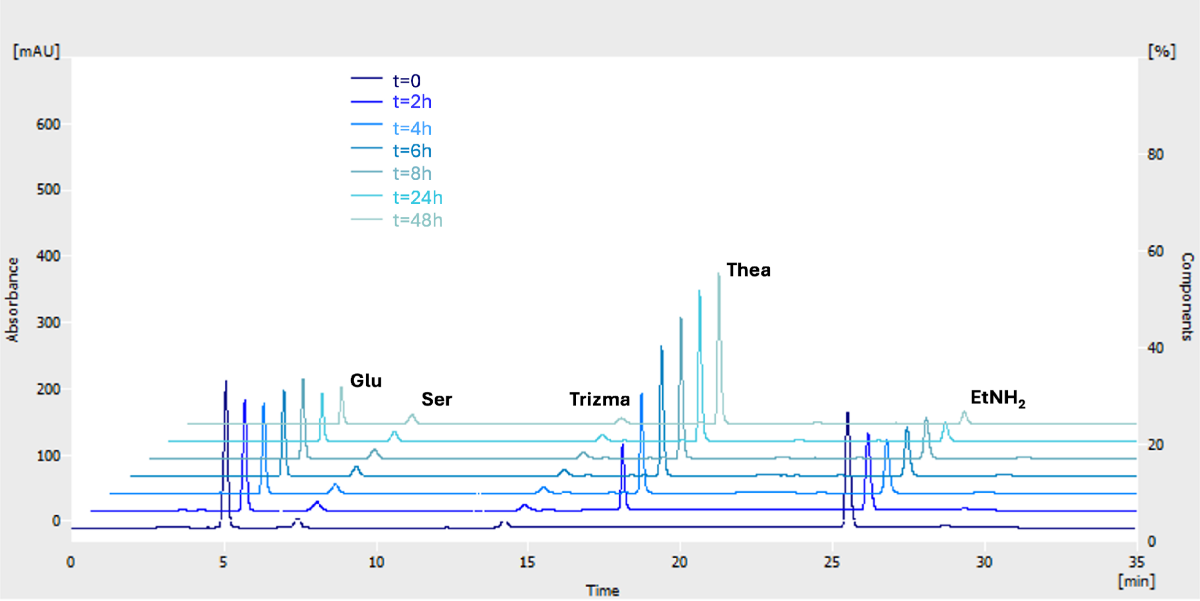

**Figure S7. HPLC chromatogram of the *in vitro* enzymatic synthesis of L-Thea**.

All samples were derivatized with OPA reagent following a standard protocol with slight modifications (Perucho J. et al., 2015). Briefly, samples (5-10 µL) were added to the derivatization mixture: 135-130 µL borate buffer (0.4 M, pH 10.4), 10 µL L-serine (2.5 mM in borate buffer) (internal standard), 50 µL OPA reagent mixture (10 mg OPA in 100 µL methanol and 900 µL borate buffer + 50 µL 3-mercaptopropionic acid). The mixture was incubated on rolling shaker at 25 °C for 2 min. The reaction was quenched by adding 5 µL of acetic acid 5% v/v and immediately analyzed by HPLC. RP-HPLC analyses were carried out on a Luna Omega Polar C18 Phenomenex column (5 µm x 150 mm x 4.6 mm) using a linear gradient of eluent A [95% sodium acetate 0.05 M pH 5.88 + 5 % v/v methanol] and eluent B (water: methanol 30 : 70): 0–2.5 min A : B 75 : 25 isocratic, 2.6–10 min A : B 67 : 33 linear gradient; 10.1–11 min A : B 40 : 60 jump; 11.1–15 min A : B 40 : 60 isocratic; 15–15.1 min A : B 20 : 80 jump; 15.1–25 min A : B 20 : 80 isocratic; 25–25.1 min A : B 75 : 25 jump; 25.1–35 min A : B 75 : 25 isocratic; flow rate: 0.5 mL/min; UV-detection: 340 nm; column: 40 °C, injection volume: 10 µL.

| time (h) | Area Thea | Area Ser | Area Thea/area Ser |
|---|---|---|---|
| 0 | 2200,18 | 211,124 | 10.42 |
| 2 | 2194,516 | 213,541 | 10.27 |
| 4 | 2223,081 | 212,832 | 10.44 |
| 6 | 2134,648 | 207,933 | 10.26 |
| 24 | 2280,348 | 207,926 | 10.96 |
| 48 | 2213,849 | 206,347 | 10.72 |
| 72 | 2256,955 | 209,798 | 10.75 |

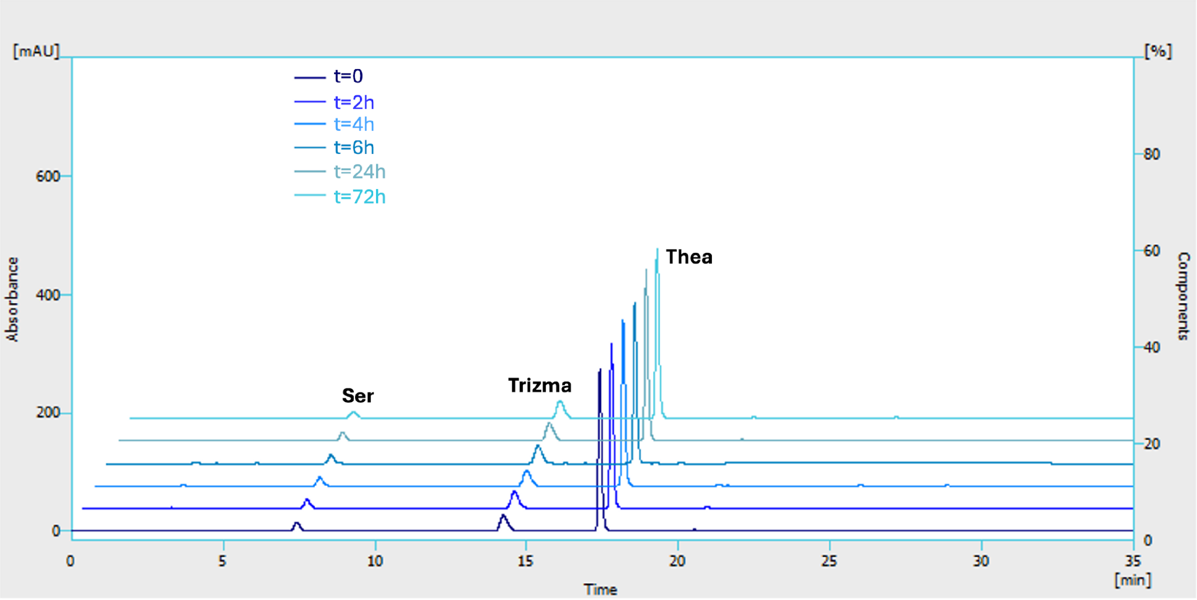

A

| time (h) | Area Thea | Area Ser | Area Thea/area Ser |
|---|---|---|---|
| 0 | 2214,248 | 204,722 | 10.81 |
| 2 | 2262,675 | 209,099 | 10.82 |
| 4 | 2305,916 | 208,809 | 11.04 |
| 6 | 2526,182 | 207,581 | 12.16 |
| 8 | 2592,971 | 209,145 | 12.39 |
| 24 | 2281,227 | 213,207 | 10.69 |
| 48 | 2385,971 | 214,204 | 11.14 |

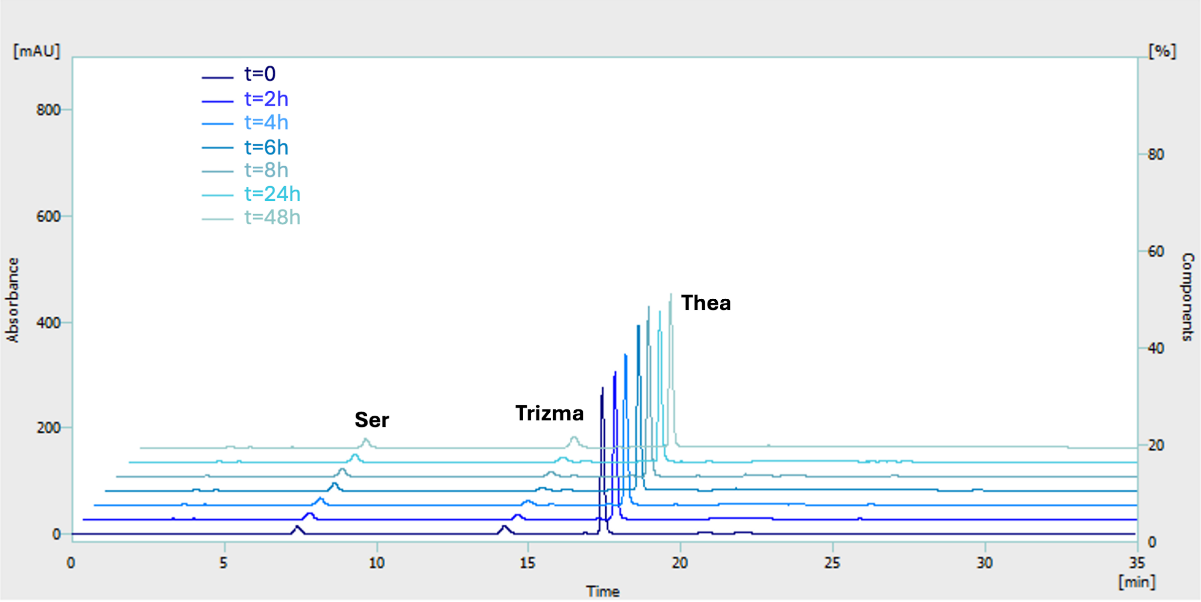

B

**Figure S8**. **HPLC chromatogram showing L-Thea stability under reaction conditions *in vitro***, in the absence (**A**) or presence of the recombinant *Mm*GMAS (**B**).

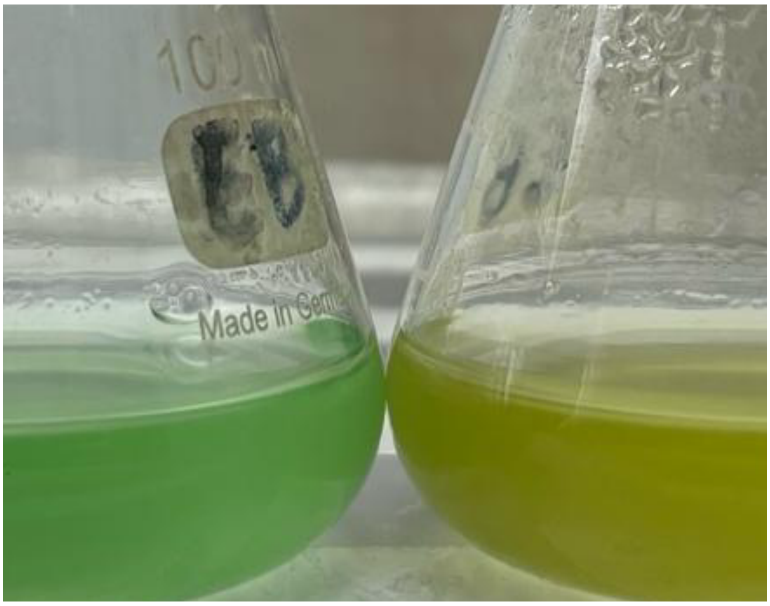

A

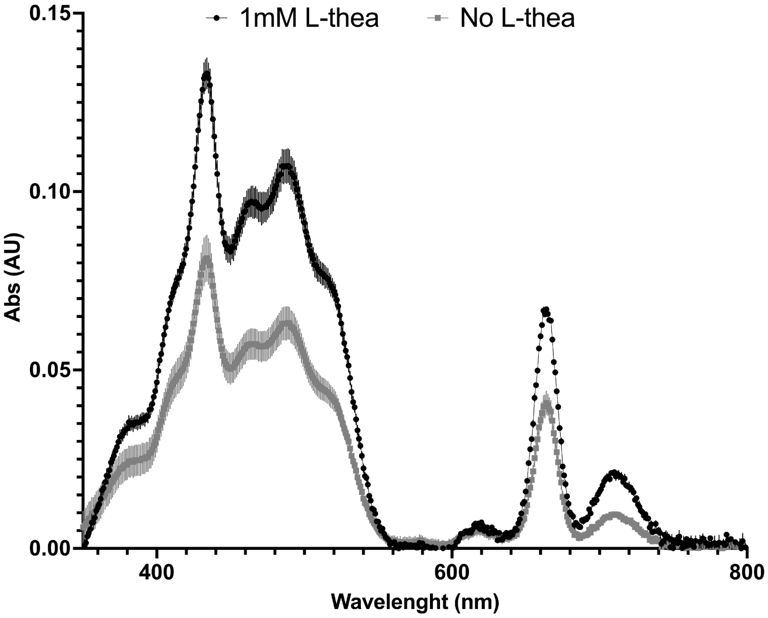

B

**Fig. S9. Effect of L-theanine in nitrogen depleted cultivation**. (A) Picture taken after 48 hours of *Synechocystis* wild-type cultivation in modified BG11. On the left, wild-type strain cultivated in modified BG11 supplemented with 1 mM **L**-theanine. On the right, same culture in the absence of L-theanine. (B) Absorption spectra of *Synechocystis* wild-type strain cultivated for 48h in modified BG11 medium both in the presence and absence of 1 mM L-theanine), and extracted in *N,N*-dimethylformamide (DMF). Cellular content was assessed prior to extraction and used for absorbance normalization. Experiment was performed in three independent replicates.

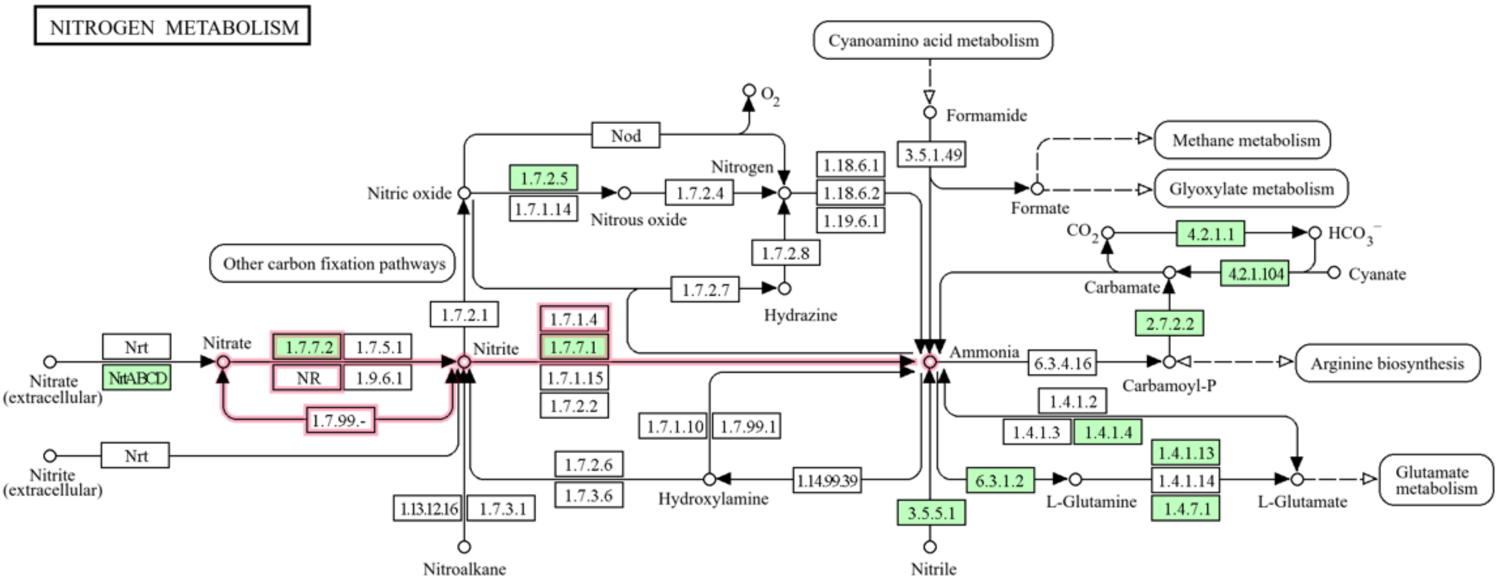

| Entry name | Locus | Activity |
|---|---|---|
| NrtABCD | *Sll*1450, *slll*1451, *sll*1452, *sll*1453, | Nitrate/nitrite transport proteins. |
| 1.7.7.2 (NarB) | *Sll*1454 | Nitrate reductase. |
| 1.7.7.1(NirA) | *Slr*0898 | Ferredoxin-nitrite reductase. |

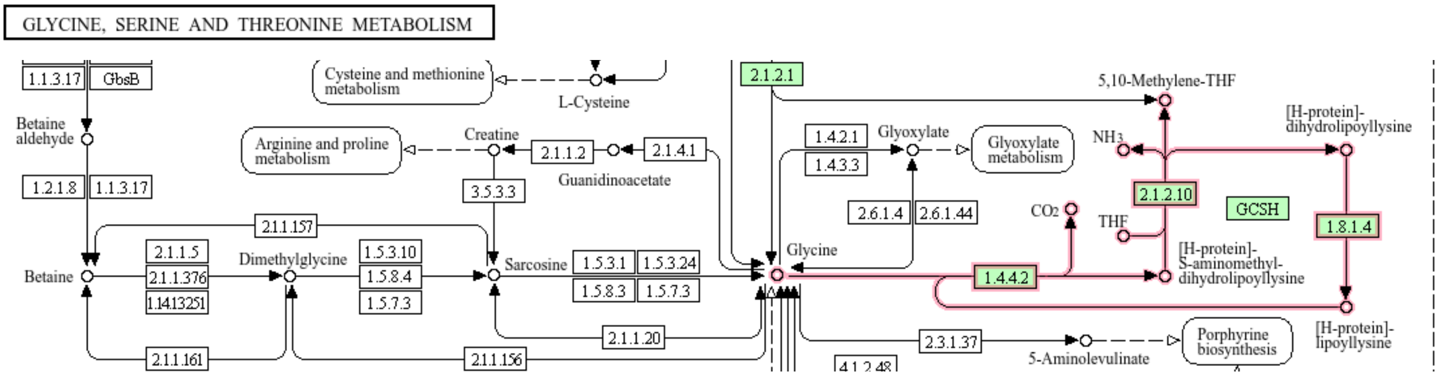

| Entry name | Locus | Activity |
|---|---|---|
| 1.4.4.2 (gcvP) | *Slr*0293 | P protein of glycine cleavage complex |
| 2.1.2.10 | *Sll*0171 | Aminomethyltransferase |

**Fig. S10. Maps of *Synechocystis* ammonia-producing metabolic pathways.** Above is the map of nitrogen metabolism in *Synechocystis* along with a related table containing name, associated loci, and descriptions of the activities of the involved enzymes. Reaction steps of interest are highlighted by red arrows, while enzymes with characterized and proven activity are highlighted in green. Below is the map of glycine, serine and threonine metabolism of *Synechocystis* along with its related table containing name, associated loci, and descriptions of the activities of the involved enzymes. The investigation, figures and tables were made using KEGG Database, specifically the KEGG Compound and KEGG Pathways databases.

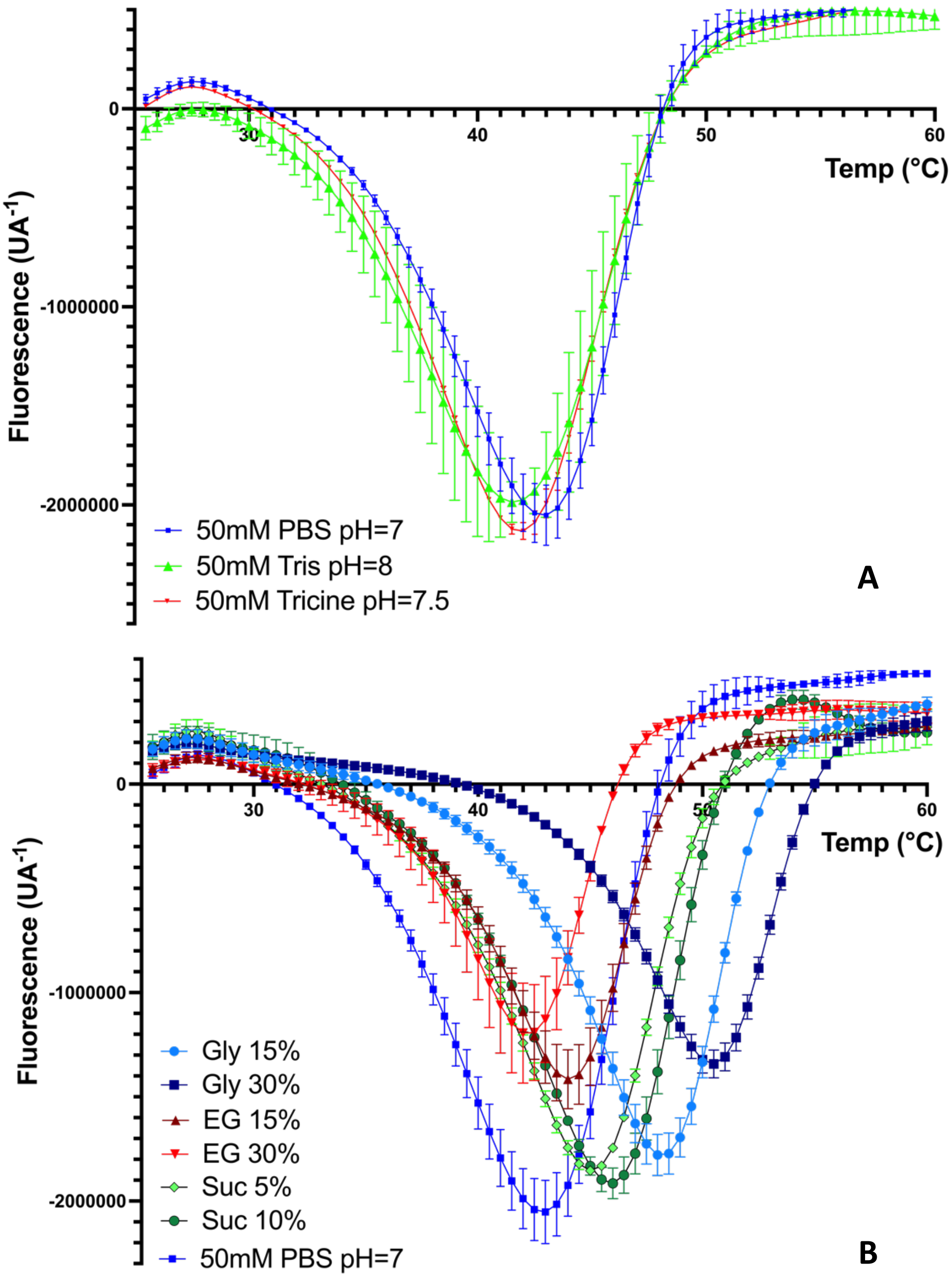

**Fig. S11. First derivatives of melting curves of recombinant *Mm*GMAS.** Derivatives melting curves calculated in the presence of (A) different buffers and (B) different additives (50mm PBS pH 7.0 +15% and 30% sucrose-w/v, ethylene glycol and glycerol-v/v) by Differential Scanning Fluorimetry (DSF). Experiments were performed in duplicate.

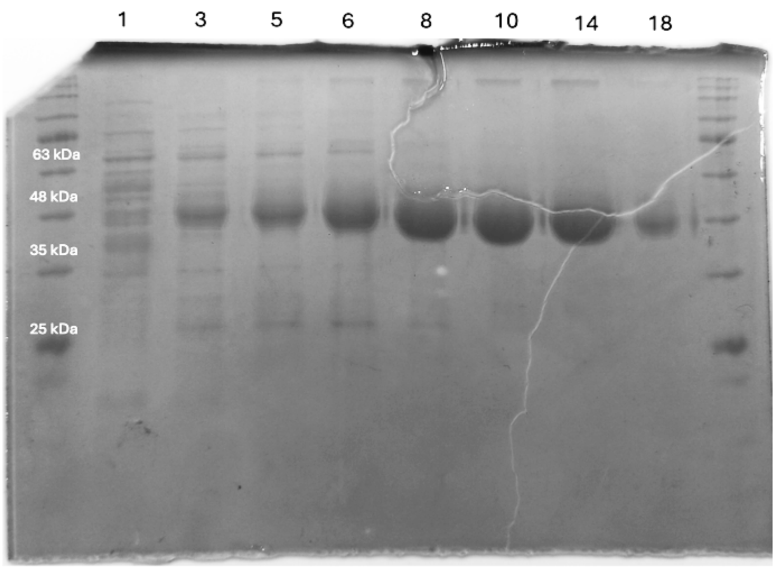

**Fig. S12. SDS-PAGE analysis of *Mm*GMAS purified under optimized conditions.** SDS-PAGE 12% acrylamide of *Mm*GMAS purification fractions collected after the IMAC purification elution gradient. Fractions from 6 to 18 were collected and used for buffer exchange, 6xHis-Tag cleavage. Prestained SharpMassTM VI protein ladder was used as protein marker (first and last lane).

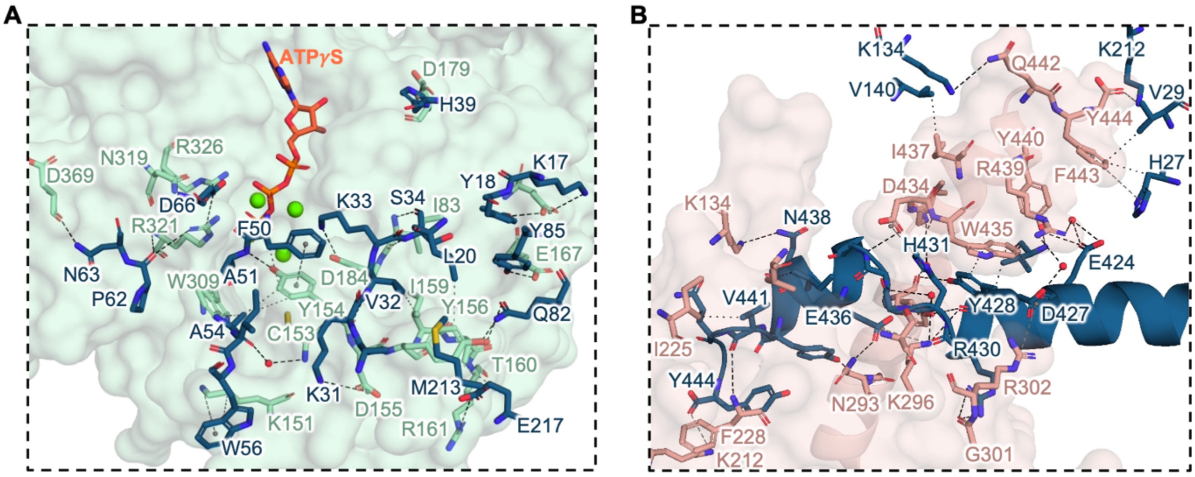

**Fig. S13. Relevant contacts between *Mm*GMAS chains.** (**A**) Structural determinants of the interaction between two adjacent *Mm*GMAS monomers belonging to the same hexamer ring. (**B**) Structural determinants of the interaction between two adjacent *Mm*GMAS monomers belonging to different hexamer rings.

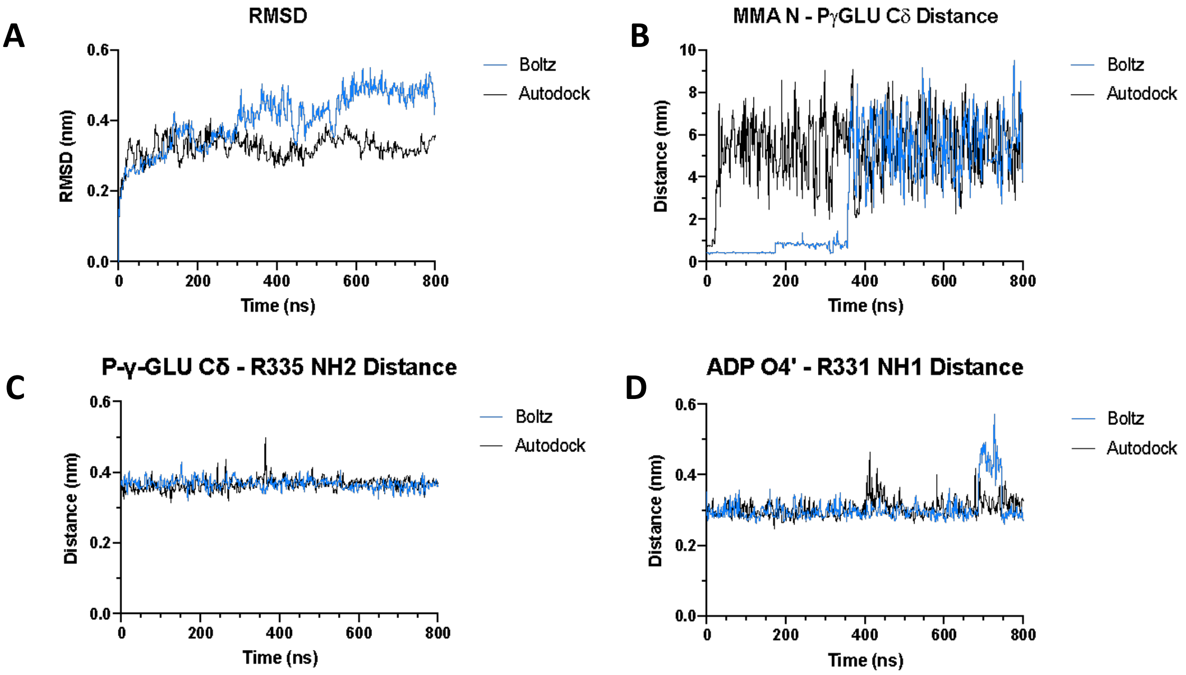

**Fig. S14. Molecular dynamics of *Mm*GMAS/ADP/P-γ-Glu/MMA.** Each panel compares MD simulations on the quaternary complex, with MMA positioned either in the binding site proposed by our work by Boltz or docked by AutoDock Vina in the binding site hypothesized by Wang et al., 2021. (**A**) RMSD analysis showing MD convergence. (**B**) Distance between the nitrogen atom of MMA and P-γ-Glu Cδ along the simulations, showing the higher stability of the MMA binding pose modelled by Boltz. **(C)** Distance between P-γ-Glu Cδ and Arg335 NH2, showing the high positional stability of P-γ-Glu along both simulations. **(D)** Distance between ADP O4' and Arg331 NH1, showing the high positional stability of ADP along both simulations.

| Name | Sequence 5’[illegible]3’ | Purpose |
|---|---|---|
| NR1 | TGGCAAATTCAAGGGGTGGT | External NS region amplification |
| NR2 | TTACCGATGAGTGGCACGTT | |
| Colony_SuperP_Rev | AGTGGGGTAGGGAAGACTGTTGC | Pcpc560 amplification |
| Colony_Kan_For | AACTCTGGCGCATCGGGCTT | Kanamycin amplification |

**Tab. S1. List of oligonucleotides used in this study**

| Medium | Composition | Purpose |
|---|---|---|
| Luria-Bertani | 10 g/L NaCl, 10 g/L Tryptone extract, 5 g/L Yeast extract. | *E. coli* cultivation |
| Standard BG11 | 10 mM N-[Tris(hydroxymethyl)methyl]-2-aminoethanesulfonic acid (TES) pH 8.0, 6 mg/L ferric ammonium citrate, 30.5 mg/L $K_2HPO_4$, 20 mg/L $Na_2CO_3$, 2.86 mg/L $H_3BO_3$, 1.81 mg/L $MnCl_2$, 0.22 mg/L $ZnSO_4$, 0.39 mg/L $NaMoO_4$, 0.08 mg/L $CuSO_4$, 0.05 mg/L $Co(NO_3)_2$, 1.49 g/L $NaNO_3$, 75 mg/L $MgSO_4$, 36 mg/L $CaCl_2$, 9.2 mg/L citric acid and 2.8 µM EDTA pH 8.0. | *Synechocystis* standard cultivation |
| Modified BG11 | 10 mM N-[Tris(hydroxymethyl)methyl]-2-aminoethanesulfonic acid (TES) pH 8.0, 6 mg/L ferric ammonium citrate, 30.5 mg/L $K_2HPO_4$, 20 mg/L $Na_2CO_3$, 2.86 mg/L $H_3BO_3$, 1.81 mg/L $MnCl_2$, 0.22 mg/L $ZnSO_4$, 0.39 mg/L $NaMoO_4$, 0.08 mg/L $CuSO_4$, 0.04 mg/L $CoCl_2$, 75 mg/L $MgSO_4$, 36 mg/L $CaCl_2$, 9.2 mg/L citric acid and 2.8 µM EDTA pH 8.0. | Nitrogen-depleted medium for *Synechocystis* cultivation |

**Tab. S2. List and composition of medium used in this study.**

| | |
|---|---|
| PDB ID | 9QUR |
| **Data collection statistics** | |
| Diffraction source | ID30A-3 (ESRF) |
| Wavelength (Å) | 0.968 |
| Temperature (K) | 100 |
| Detector | Eiger1_4M |
| Crystal-detector distance (mm) | 214.28 |
| Rotation range per image (°) | 0.2 |
| Exposure time per image (s) | 0.01 |
| Space group | C 1 2 1 |
| No. of molecules/ASU | 6 |
| a, b, c (Å) | 141.740 246.923 105.760 |
| α, β, γ€(°) | 90.00 111.17 90.00 |
| Total no. of reflections | 595255 (25068) |
| No. of unique reflections | 97025 (4525) |
| Completeness (%) | 98.7 (93.0) |
| Redundancy | 6.1 (5.5) |
| ⟨I/σ(I)⟩ | 4.1 (0.9) |
| Rmrg | 0.365 (1.310) |
| **Refinement statistics** | |
| Resolution range (Å) | 90.89 - 2.65 |
| No. of reflections, working set | 94983 |
| No. of reflections, test set | 2031 |
| Final Rcryst | 0.225 |
| Final Rfree | 0.259 |
| No. of non-H atoms | |
| Protein | 20738 |
| $Mg^{2+}$ ions | 18 |
| ATPγS | 186 |
| Water | 103 |
| Total | 21045 |
| R.m.s. deviations | |
| Bonds (Å) | 0.0055 |
| Angles (°) | 1.435 |
| Average B factors ($Å^2$) | 37.0 |
| Ramachandran plot | |
| Most favored (%) | 97% |
| Allowed (%) | 3% |
| RSCC ATPγS (Chain ID) | 0.87 (F), 0.88 (C, A), 0.89 (B), 0.90 (E, D) |

**Tab. S3. Crystallographic data statistics**
